# No genome-wide correlations and little shared genetic architecture between reproductive life-history traits and estrogen receptor-positive breast cancer risk

**DOI:** 10.1101/2025.06.26.661731

**Authors:** Euan A. Young, E Yagmur Erten, Erik Postma, Virpi Lummaa, Bert van der Vegt, Geertruida H de Bock, Hannah L. Dugdale

## Abstract

Estrogen receptor-positive breast cancer (ER+ BC) is one of the most prevalent cancers, but the evolutionary processes shaping genetic variation in ER+ BC risk are poorly understood. Both evolutionary life-history theory and evidence from studies of individual ER+ BC risk variants suggest that increased genetic ER+ BC risk is associated with faster maturation, earlier reproduction, and/or increased reproductive success (i.e., there is a trade-off), but it is unclear how well this pattern is replicated when considering the polygenic architecture of these traits after controlling for potential biases. Here, we estimate genome-wide genetic correlations between ER+ BC risk and three reproductive traits (age at menarche, age at first birth, and the number of children) using genomic restricted maximum-likelihood analyses on Lifelines biobank data and linkage disequilibrium score regressions on population and family-based genome-wide association study data. Regardless of the data or method used, genetic correlations were low and not statistically significant. Further analyses decomposing genome-wide genetic variance into local regions detected only three loci exhibiting significant pleiotropy between ER+ BC risk and age at menarche, suggesting little shared genetic architecture between ER+ BC risk and reproductive traits. Thus, the role of life-history trade-offs in shaping ER+ BC risk in European populations appears, at most, small, and the evolutionary processes giving rise to this life-threatening disease remain unclear. Future studies could examine the impact of evolutionary mismatches in shaping ER+ BC risk, where conducting longitudinal studies on populations transitioning to reproductive patterns observed in contemporary European populations would be most useful.

## INTRODUCTION

Breast cancer is the fourth most deadly cancer worldwide (GBD, 2021) and accounts for the greatest proportion of new cancer cases (Sung et al., 2021). Estrogen receptor-positive breast cancer (ER+ BC) is the most common subtype, accounting for ∼80% of all breast cancer cases overall, and is most common after age ∼50 (Yasui & Potter, 1999). In European ancestry populations, common germline genetic variation explains ∼14% of the variation in a woman’s risk of ER+ BC (Zhang et al., 2020), and over 200 genome-wide significant risk variants have now been identified (Michailidou et al., 2013, 2017; Zhang et al., 2020). However, we have a poor understanding of why genetic variation promoting a life-threatening disease persists in contemporary populations.

Cancer is common not only in humans, but across the tree of life (Aktipis, Boddy, et al., 2015; Boddy et al., 2015), and organisms have been under selective pressure to evolve cancer suppression mechanisms since the dawn of multicellularity over half a billion years ago (Merlo et al., 2006; Nunney, 2013). Evolutionary biology hypothesizes that genetic risk variants for diseases such as cancer may persist in populations due to their beneficial effects at younger ages i.e., because of life-history trade-offs (*life history theory* and the *antagonistic pleiotropy theory of aging*, Jacqueline et al., 2017; Promislow & Harvey, 1990; Roff, 1992; Stearns, 1977, 1992; Williams, 1957). For example, genetic variants promoting a faster life-history (earlier maturation and reproduction) could be selected for if they result in higher reproductive success, even if they increase the risk of cancer in later life (Figure 1, Dujon et al., 2022; Jacqueline et al., 2017). Altogether, these two non-mutually exclusive theories could explain why variation in the strength of cancer suppression covaries with life history, such as why cancer prevalence is higher in bird species with larger clutch sizes (Kapsetaki et al., 2024). Thus, due to reproductive traits’ close association with fitness, examining how genetic variation in ER+ BC risk is associated with reproductive traits could help us understand the evolutionary processes shaping ER+ BC risk variants, including those under contemporary directional selection (Haldane, 1954; Lande & Arnold, 1983; Mathieson et al., 2023; Robertson, 1966).

**Figure 1:**
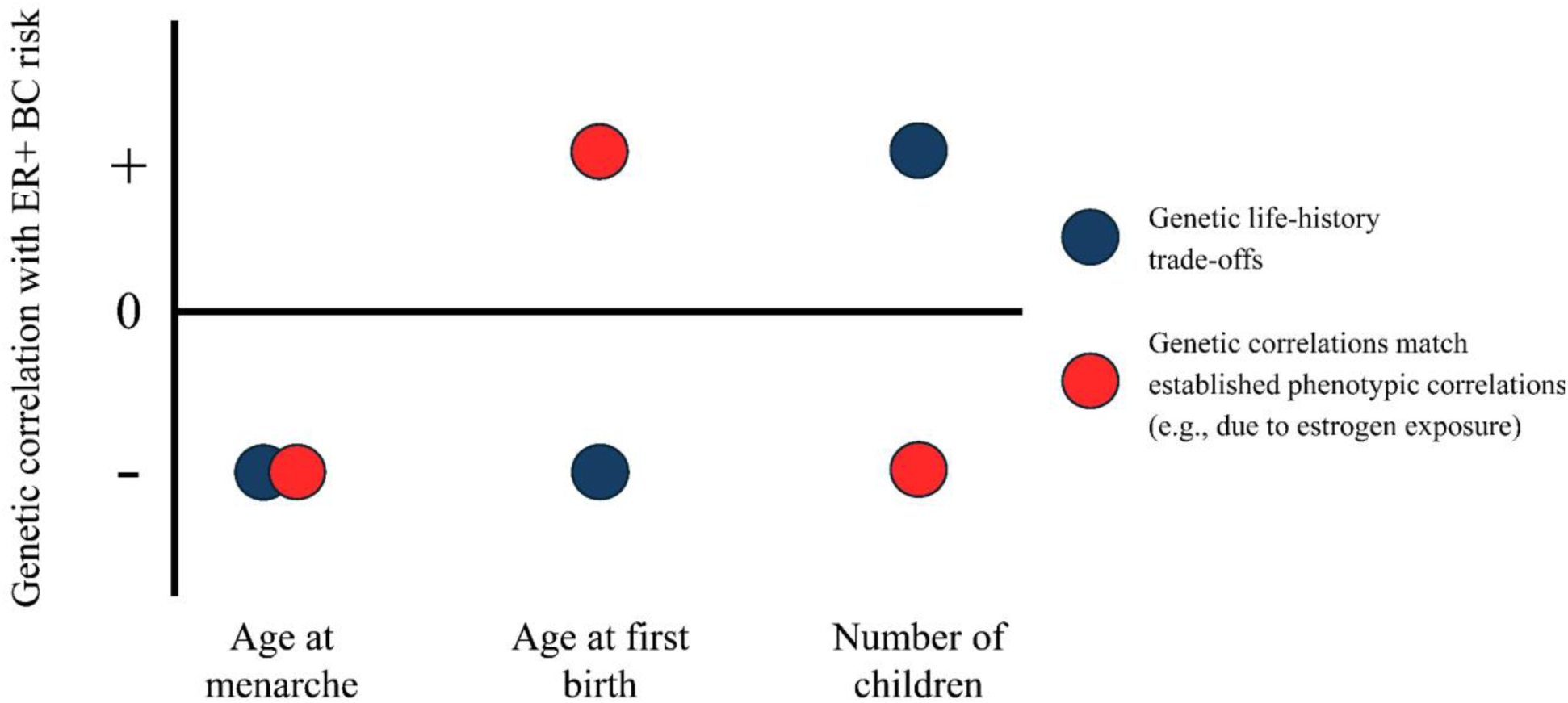
Predicted direction of genetic correlations between ER+ BC risk and reproductive life-history traits (age at menarche, age at first birth, and number of children) under different hypotheses. Circles below the x-axis represent negative genetic correlations, i.e., genetic predispositions for lower/earlier values for the life-history trait are correlated with increased ER+ BC risk. Circles above the x-axis represent positive genetic correlations, i.e. genetic predispositions for higher/later values for the reproductive trait are correlated with increased ER+ BC risk. Circle colors indicate different hypotheses: blue, *genetic life-history trade-offs* as predicted by antagonistic pleiotropy theory of ageing and life-history theory; and red, *genetic correlations match established phenotypic correlations (e.g. due to estrogen exposure)*, where genetic correlations match the existing evidence from phenotypic studies of how life-history traits shape ER+ BC risk phenotypically, potentially due to their influence on a woman’s lifetime estrogen exposure.

In line with these evolutionary theories, individuals with genetic predispositions to earlier maturation, reproduction, and/or higher numbers of children also show faster aging (Long & Zhang, 2023; Wang et al., 2013) and increased risk for specific diseases (Byars et al., 2017; Perry et al., 2015; Trumble et al., 2023). Cancer appears no exception, with a host of tumor suppressor genes being linked to the regulation of puberty and fertility (reviewed in Boddy et al., 2015; Byars & Voskarides, 2020; Crespi & Summers, 2006; Dujon et al., 2022; Jacqueline et al., 2017). These include *p53* genetic variants increasing conception likelihoods and overall cancer risk (Kang et al., 2009), and shorter repeats of the *CAG* repeat region of the androgen receptor promoting both increased fertility and prostate cancer risk (Summers & Crespi, 2008). Breast cancer (and in particular ER+ BC) is a good candidate for the maintenance of disease risk variants by life-history trade-offs, considering that genetic variation in breast cancer is enriched in gene-expression hubs involved in estrogen metabolism and signaling (Stone et al., 2025), and that these hormones play central roles in both fertility (through regulating processes such as conception, implantation, and pregnancy) as well as the etiology of ER+ BC (Ghosh et al., 2020; Jacqueline et al., 2017; Jasienska et al., 2017). Indeed, one genealogical study from Utah found that carriers of BRCA1/2 mutations also had more children (Smith et al., 2012), and contemporary studies examining all genome-wide significant loci for age at menarche found that genetic variants lowering age at menarche were also associated with higher overall breast cancer risk (Xiang et al., 2025) and, specifically, higher ER+ BC risk (Day et al., 2017; Zhao et al., 2024).

Contrary to these genetic studies, phenotypic associations from epidemiological studies are not entirely consistent with the predictions from evolutionary life-history theory (Figure 1), potentially owing to the influence of an evolutionary mismatch. In addition to life-history trade-offs, evolutionary mismatches are also important in cancer evolution, where populations experiencing rapid cultural, environmental, and/or lifestyle changes may experience an evolutionary mismatch between their life history and evolved cancer suppression mechanisms (Aktipis & Nesse, 2013; Nesse, 2011, 2019). Relevant for ER+ BC in contemporary European populations are changes in diet and nutrition which have caused earlier ages at menarche (Duan et al., 2020; Nguyen et al., 2020; Rees, 1993) and recent changes in cultural preferences towards smaller families driving later ages at first birth and lower numbers of children (Aitken, 2022; Kirk, 1996). Because amenorrhea (an absence of menstruation) is induced during breastfeeding, both lower family sizes and earlier ages at menarche may lead to women experiencing more menstrual cycles and a higher lifetime exposure to estrogen than women’s cancer suppression mechanisms have evolved to handle (Coe & Steadman, 1995; Eaton et al., 1994; Short, 1976; Strassmann, 1999) due to estrogen’s proliferative effects on mammary cells (Dall et al., 2018). Phenotypic evidence supports this theory and, while earlier ages at menarche have been associated phenotypically with higher ER+ BC risk (in line with evolutionary life-history trade-offs), earlier ages at first birth and higher numbers of children are actually associated with lower ER+ BC risk (Figure 1, Aktipis et al., 2015; Kelsey et al., 1993; Momenimovahed & Salehiniya, 2019), potentially owing to their correlation with lifetime estrogen exposure (Aktipis, Ellis, et al., 2015; Bieuville et al., 2023). Consequently, how genetic predispositions to ER+ BC are associated with reproductive traits may deviate from predictions made by life-history theory (Figure 1) and remain unclear.

A complete understanding of how genetic predispositions to ER+ BC are associated with reproductive traits in contemporary European populations is currently lacking due to – first and foremost – the importance of non-genetic factors on both ER+ BC risk and reproductive traits. Because ER+ BC risk and reproductive traits are influenced by both lifestyle factors and socioeconomic and cultural backgrounds (Barban et al., 2016; Colditz et al., 2004), separating these environmental factors from genetic associations is crucial for unmasking any evolutionary processes, including life-history trade-offs or genetic factors that underlie documented phenotypic associations. Second, both ER+ BC risk and reproductive traits are highly polygenic Hence, previous genetic studies focusing on genome-wide significant variants fail to capture the full scope of genetic variation in both ER+ BC risk and reproductive traits. Indeed, genome-wide significant variants explain less than 50% of the total variance in genomic risk for ER+ BC (Zhang et al., 2020), and only 25% of the total genetic variation in age at menarche (Day et al., 2017). Even fewer genome-wide significant variants for age at first birth and number of children born have been detected, despite significant heritability for both traits (Barban et al., 2016; Mathieson et al., 2023; Mills et al., 2021; Tropf et al., 2015).

To resolve this knowledge gap, we need to quantify the overall extent and effects of shared genetic architecture between ER+ BC risk and key reproductive traits: age at menarche, age at first birth, and number of children. First, genome-wide genetic correlations – which estimate the aggregated contribution of pleiotropy and linkage disequilibrium (LD) to the covariation of traits (van Rheenen et al., 2019) – can be used to measure how genetic factors shape observed phenotypic associations, on average, across all loci. To the best of our knowledge, studies estimating genome-wide genetic correlations between ER+ BC risk and human life-history traits are limited to two studies estimating genetic correlations between ER+ BC risk and age at menarche (Jiang et al., 2019; Prince et al., 2024), one of which also estimated the genetic correlations between other life-history traits, including age at first birth and number of children (Prince et al., 2024). These studies revealed negligible genetic correlations between all life-history traits and ER+ BC risk, except for a small (r_g_=-0.048) and statistically non-significant (p = 0.062) genetic correlation between the number of children and ER+ BC risk. However, these studies did not fully account for confounding factors such as population stratification, which can bias genetic correlations (Berg et al., 2019; Border et al., 2022), particularly those involving reproductive traits (Howe et al., 2022; Tan et al., 2024). Second, because genome-wide genetic correlations estimate *average* genetic associations across all loci, they can miss smaller regions of shared genetic architecture that may have important influences on traits (Shi et al., 2017; van Rheenen et al., 2019). Therefore, it is also necessary to examine regional heritability and genetic correlations to understand the extent to which genetic predispositions to ER+ BC are shaped by human reproductive traits.

Here we explore evidence for human reproductive traits shaping genetic predispositions to ER+ BC, accounting for environmental confounding and the polygenicity of these traits. We first estimate genome-wide heritabilities and genetic correlations by performing a genomic restricted maximum-likelihood (GREML) analysis (Lee et al., 2012; Yang et al., 2010) using the Dutch biobank Lifelines (Scholtens et al., 2015) linked to national pathology records (Casparie et al., 2007). We then perform linkage disequilibrium (LD) score regressions (LDSC, Bulik-Sullivan et al., 2015) on the largest available genome-wide association study (GWAS) data to estimate genetic correlations at the population level across all available loci (i.e., population effect GWAS data; Loh et al., 2018; Mathieson et al., 2023; Michailidou et al., 2017; Mills et al., 2021). Subsequently – and owing to the complex influence of environmental and cultural factors upon these traits – we estimate genetic correlations from GWAS data using family-based approaches (i.e., direct genetic effect GWAS data; Tan et al., 2024) which exploit the random Mendelian segregation among siblings to remove environmental confounding from genetic correlations due to factors such as population stratification and assortative mating (Kong et al., 2018; Young et al., 2022; Young, 2024). Finally, we divide the genome into 1,703 approximately LD-independent genomic regions (Berisa & Pickrell, 2016) and estimate the contribution of each of these loci to the total heritability of each trait and the genetic covariance between ER+ BC risk and reproductive traits (*ρ*-HESS, Shi et al., 2017). Together, these approaches allow us to quantify the role genetic variation plays in explaining phenotypic associations between ER+ BC risk and reproductive traits overall and evaluate the extent of shared genetic architecture and pleiotropy in our search for the evolutionary processes giving rise to a life-threatening disease.

## RESULTS

### GREML analyses of Lifelines data

We first performed GREML analyses on data from the Lifelines biobank (*n* = 97,131 women) which leverages variation in genomic relatedness among non-relatives to derive heritability estimates and genetic correlations. Using the largest available genomic dataset in Lifelines (Global screening array, *n* = 535,107 SNPs), univariate analyses revealed statistically significant heritability (±SE) for age at menarche (43.1±5.8%, *n* = 8,071 women) and age at first birth (24.2±7.4%, *n* = 6,260 women), suggesting that genetic variance plays a significant role in shaping these traits (p < 0.05, Table 1a). The heritability of the number of children was estimated with less precision and was not statistically significant (12.8±9.8%, p = 0.337, *n* = 4,602 women, Table 1a). This was also true for ER+ BC risk, where the heritability was estimated as 11.8% (±5.2%, p = 0.190, *n* = 182 cases and *n* = 8,700 controls, Table 1a).

**Table 1:** Heritability estimates (and standard error, SE) of traits from **a)** GREML analyses using Lifelines genotype data, **b)** LD score regression on population effect and **c)** direct genetic effect GWAS data. P-values from likelihood ratio tests (LRT, for GREML) and z-scores (for LD score regressions) and results are presented in bold when significant (< 0.05). Results from the Lifelines Global screening array data are shown. Results from analyses of other Lifelines genotype data are shown in Table S1.

| Trait | a) GREML on Lifelines data |  | b) LD score regressions on population effect GWAS data |  | c) LD score regressions on direct genetic effect GWAS data |  |
| --- | --- | --- | --- | --- | --- | --- |
| | Heritability±SE | P-value | Heritability±SE | P-value | Heritability±SE | P-value |
| ER+ BC risk | 11.8±5.2% | 0.190 | **10.0±1.0%** | **< 0.001** | - | - |
| Age at menarche | **43.1±5.8%** | **< 0.001** | **20.0±0.8%** | **< 0.001** | **9.3±2.5%** | **< 0.001** |
| Age at first reproduction | **24.2±7.4%** | **< 0.001** | **4.3±0.8%** | **< 0.001** | **3.2±1.0%** | **< 0.001** |
| Number of children | 12.8±9.8% | 0.337 | **2.2±0.1%** | **< 0.001** | **2.7±0.8%** | **< 0.001** |

These heritability estimates were largely consistent when using other types of genomic data available in Lifelines (the Affymetrix, *n* = 441,001 SNPs, and CytoSNP genomic array data, *n* = 244,041 SNPs, Table S1). However, the heritability estimate for ER+ BC risk became statistically significant when using the Affymetrix array data (p = 0.04, Table S1), likely because these data contained a slightly higher number of individuals (*n* = 198 cases, *n* = 8,972 controls). Moreover, the heritability estimate for age at menarche was considerably lower (but remained statistically significant) when using the CytoSNP data (17.0% ±5.2%, p < 0.001, *n* = 6,155 women, Table S1), most likely because of the lower genomic coverage of the CytoSNP data (Table S2).

Genetic correlations (±SE) between ER+ BC risk and life-history traits were all low and statistically non-significant when using the Lifelines data (p > 0.05/3, to correct for multiple testing across reproductive traits, Figure 2, purple, Table S3). Using the global screening array data these genetic correlations were 0.05±0.17 for age at menarche (*n* = 176 cases, *n* = 7,895 controls), 0.10±0.24 for age at first birth (*n* = 155 cases, *n* = 6,105 controls), and 0.10±0.35 for number of children (n = 146 cases, n = 4,456 controls, p > 0.016, Fig 2a, Table S2). Precision of these estimates was also low, but a power analysis estimated that we had 80% power to detect genetic correlations between ER+BC risk and the life-history trait stronger than ±0.29 for age at menarche, ±0.40 for age at first birth, and ±0.52 for the number of children (Figure S1). These null findings were replicated when using both the Affymetrix and CytoSNP data (Table S3), with slightly reduced power in the CytoSNP data (Figure S1), which was expected given its lower sample size.

**Figure 2:**
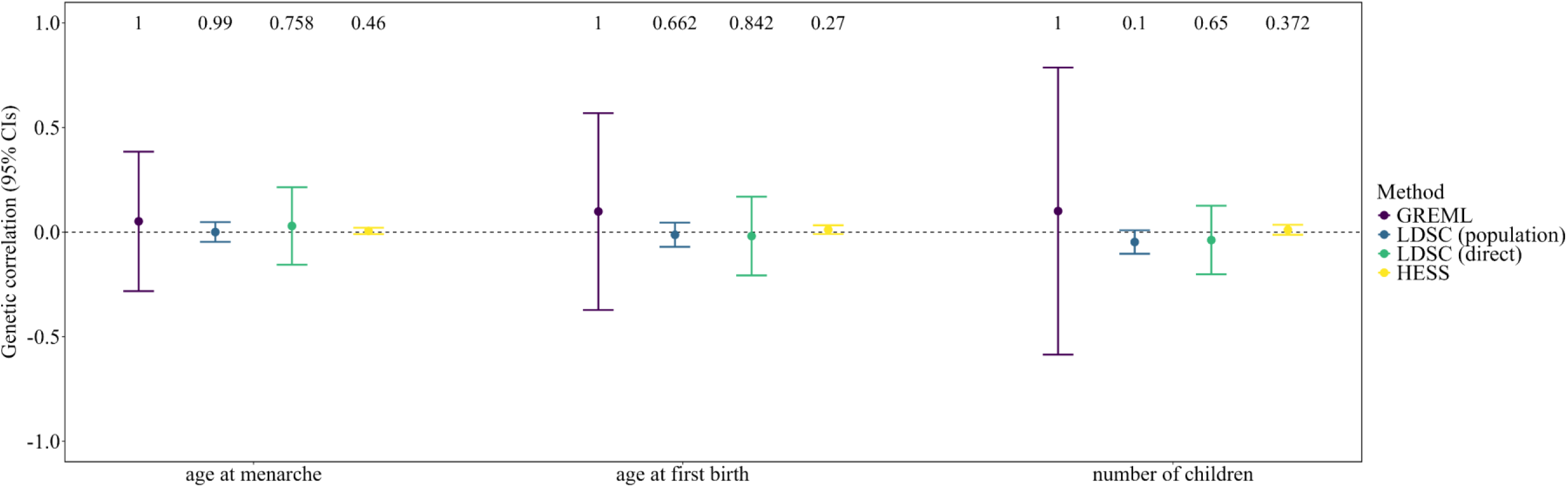
Global genetic correlations between ER+ BC risk and three reproductive traits: age at menarche, age at first birth, and number of children. Legend and colors indicate which analysis was used to estimate genetic correlations: genomic relatedness matrix (GREML) analyses using Lifelines biobank data (purple); LD score regression analyses (LDSC) on population (blue) and direct genetic effects (green), and HESS analyses (yellow), all using GWAS summary statistics. Circles above the dotted line indicate positive genetic correlations, and circles below indicate negative genetic correlations. Error bars show 95% confidence intervals approximated from SEs. P-values testing for a non-zero genetic correlation are shown above each trait. For GREML analyses, these were derived from LRTs. For LDSC and HESS analyses, these were approximated from z-scores. Significance was adjusted for multiple testing across three traits (p < 0.05/3).

### LD score regressions of GWAS data

We then performed higher-powered analyses using the largest available GWAS data on European-ancestry individuals for all traits with LD score regressions (ER+ BC risk, *n* = 11,792,542 SNPs, *n* = 69,501 cases and *n* = 105,974 controls, Michailidou et al., 2017; for age at menarche, *n* = 11,984,352 SNPs, *n* = 242,278, Loh et al., 2018; for age at first birth, *n* = 9,744,772 SNPs, *n* = 542,901, Mills et al., 2021; and for number of children, *n* = 11,792,542 SNPs, *n* = 785,604, Mathieson et al., 2023). LD score regressions estimate heritabilities and genetic correlations by regressing the genome-wide associations between traits and genetic variants available with GWAS data on their LD scores, relying on the statistical corrections undertaken by GWAS analyses to account for factors such as population stratification that may bias GWAS estimates (Bulik-Sullivan, Loh, et al., 2015). Albeit estimated with more precision and becoming statistically significant, our heritability estimate (±SE) from GWAS data for ER+ BC risk was consistent with our results from the Lifelines biobank for ER+ BC risk (10.0±1.0%, p < 0.001, *n* = 1,117,551 SNPs, Table 1a). Heritability estimates from this GWAS data for reproductive traits were also more precise but also generally decreased from estimates using GREML, likely owing to better removal of inflation owing to gene-environment correlations and population stratification (age at menarche *h*^2^ = 20.0±0.8%, *n* = 1,185,955 SNPs; age at first birth *h*^2^ = 4.3±0.8%, *n* = 1,179,108 SNPs; and number of children *h*^2^ = 2.2±0.1%, *n* = 1,183,820 SNPs; all p-values <0.001, Table 1b).

LD score regressions also provide additional summary statistics that allow inferences on the degree of inflation in GWAS data, overall, and whether this is due to high degrees of polygenicity or non-direct genetic factors, such as population stratification (Bulik-Sullivan, Loh, et al., 2015). These statistics suggest that the GWAS data from reproductive traits are somewhat inflated, especially for age at menarche (age at menarche, λ_GC_ = 1.69, intercept±SE = 1.13±0.01; age at first birth λ_GC_ = 1.47, intercept = 1.09±0.01; and number of children, λ_GC_ = 1.35, intercept, 1.06±0.01, Table S4), but, overall, the degree of inflation in the summary statistics due to factors other than polygenicity appeared low (ratio±SE, age at menarche 0.11±0.01, age at first birth = 0.14±0.02, and number of children 0.13±0.02).

To examine whether these potential biases of GWAS summary statistics were attributable to factors other than polygenicity and whether they affected estimates of heritability and genetic correlations, we ran additional LD score regressions on *direct genetic* effect GWAS data from family-based approaches (Tan et al., 2024, age at menarche, *n* effective sample size = 19,678, *n =* 5,671,230 SNPs, age at first birth = 35,982, *n =* 6,451,866 SNPs, and number of children = 41,589, *n =* 5,333,984 SNPs). These methods additionally account for parental genotypes in the estimation of GWAS data, allowing the separation of population stratification and other forms of environmental confounds from statistical associations, and essentially exploit the random segregation of alleles among siblings to estimate direct genetic effects (Kong et al., 2018; Young et al., 2022; Young, 2024). Consequently, heritability estimates decreased further for age at menarche (9.3±2.5%, *n* = 573,863 SNPs), although remained similar for age at first birth and number of children (3.2±1.0%, *n* = 703,396 SNPs, and 2.7±0.8%, *n* = 665,780 SNPs, respectively, p-values all < 0.001, Table 1c). Unfortunately, direct genetic effect GWAS data for ER+ BC risk were unavailable. However, the degree of inflation for ER+ BC risk was lower than for reproductive traits, making it less susceptible to biased estimates (λ_GC_ = 1.28), and this inflation appeared to be mostly due to polygenicity (intercept = 1.13±0.01, ratio = 0.24±0.02, Table S4).

Despite higher power and precision, estimating genetic correlations between ER+ BC risk and reproductive traits across all GWAS data revealed no statistically significant genome-wide genetic correlations (p > 0.05/3, Figure 2, blue and green, Tables S5 and S6). Direct genetic effect GWAS data produced lower precision estimates and, in some cases, estimates varied quantitatively from those based on population effect GWAS data, but overall estimates remained very low and statistically non-significant regardless of the traits’ heritabilities and type of GWAS data. Specifically, estimated genetic correlations (±SE) between ER+ BC risk and our life-history traits were: for age at menarche, using population effects = 0.000 (±0.024, p = 0.990, *n* = 1,111,653 SNPs), and using direct genetic effects = 0.029 (±0.094, p = 0.758, *n* = 551,453 SNPs); for age at first birth, using population effects = -0.013 (±0.029, p = 0.662, *n* = 1,113,353 SNPs) and using direct genetic effects = -0.019 (±0.095, p = 0.842, *n* = 668,389 SNPs); and the number of children ever born, using population effects =-0.047 (±0.029, p = 0.842, p = 0.100, *n* = 1,114,830 SNPs) and using direct genetic effects = -0.038 (±0.083, p = 0.650, *n* = 635,154 SNPs, Tables S5 and S6). In light of these results, and given their lower power, precision, and ability to remove potential biases, we recommend caution when interpreting the genetic correlations from our GREML analyses with our LD score regressions, suggesting that the genetic correlations between ER+ BC risk and reproductive traits are unlikely to be as strong as the confidence intervals from the GREML analyses imply.

We also found two statistically significant genetic correlations (p < 0.05/3) between the reproductive traits themselves using population effect GWAS data (age at first birth ∼ age at menarche = 0.097±0.021, p < 0.001 *n* = 1178825 SNPs; age at first birth ∼ number of children = 0.705±0.021, p < 0.001, *n* = 1179108 SNPs, Table S7). No significant correlations were found when using direct genetic effect estimates, but these had reduced power and directions of correlations remained similar (age at first birth ∼ age at menarche = 0.204±0.216, p = 0.344, *n* = 435,982 SNPs; rage at first birth ∼ number of children = -0.302±0.210, p = 0.151, *n* = 651,960 SNPs, Table S8).

### HESS analyses of GWAS data

Our genome-wide genetic correlations estimated by GREML and LD score regressions describe the aggregated associations across variants on ER+ BC risk and reproductive traits. Thus, while there is no indication of a genome-wide genetic correlation between these traits, pleiotropic loci associated with these traits may still be present (Shi et al., 2017). We therefore employed Heritability Estimation from Summary Statistics (*ρ-*HESS, Shi et al., 2017), which uses population-specific LD patterns to partition the genome-wide heritability and genetic correlations into 1,703 approximately LD-independent blocks for Europeans (on average 1.6Mb wide (Berisa & Pickrell, 2016)) and measure the local heritability and genetic correlations due to genetic variation across SNPs restricted to these genomic regions (Shi et al., 2017). This allows us to search for regions of significant local heritability and pleiotropic loci missed in our global analyses, thus giving a finer-scale understanding of the (shared) genetic architecture of these traits. Because it is recommended to only apply HESS to GWAS data with sample sizes greater than 50,000 (Shi et al., 2017), we used only the population effect GWAS data for these analyses.

Our analysis detected 30 loci of significant local heritability for ER+ BC risk and 152 loci for age at menarche (p < 0.05/1,703 = 2.94E-05 to correct for multiple testing across loci; Tables S9 and S10), amounting to 18.0% and 26.5% of the total estimated heritability for these traits, respectively. At most, for ER+ BC risk, one locus contributed 3.1% of the total heritability (chromosome 10:123,231,465-123,900,544), which is situated near the gene *FGFR2* previously implicated in ER+ BC risk studies (e.g., Jia et al., 2024). For age at menarche, one locus contributed at most 1.1% of the total heritability (chromosome 6:103,983,395-106,056,732). This locus harbors the gene *LIN28B* previously identified as the largest signal for age at menarche and which is thought to play a role in energy homeostasis and growth (Perry et al., 2014). For age at first birth, only two neighboring loci reached statistical significance (chromosome 3:47,727,212-49,316,971 and 49,316,972-51,832,014), and none were detected for number of children, agreeing with our previous analyses of the low heritability and highly polygenic nature of these traits. The regions of significant local heritability for age at first birth amounted to 1.4% of the total heritability and likely reflect the most significant locus in GWAS studies for age at first birth (chromosome 3:49,878,611), where genetic variants affect the genes *ACTL11P* and *MST1R* (Mills et al., 2021).

HESS further quantified the genetic covariance between reproductive traits and ER+ BC risk due to genetic variation restricted to these genomic regions and identified three regions of significant local genetic covariance between ER+ BC risk and age at menarche (p < 0.05/1,703, Figure 3, Table 2), but not for any other reproductive traits (p > 0.05/1,703, Supplementary data). The first locus (chromosome 8:76,456,542-79,132,860) positively covaried with both traits and harbors the gene *ZFHX4,* which has previously also been identified as a candidate gene affecting ages at menarche (Perry et al., 2014). For ER+ BC risk, this region did not reach statistical significance for local heritability, but an adjacent locus did (chromosome 8: 75,445,064-76,456,541, p = 2.90E-12, Table 2). Thus, this significant genetic covariance may partially reflect some genetic linkage with genetic variants found in the neighboring locus, which harbors the genes *HNF4G* and *RNU2-54P* previously associated with ER+ BC risk (Wu et al., 2024). The second locus showed negative local covariance (chromosome 16:53,382,572-55,903,773), reached statistical significance for local heritability for age at menarche (Figure 3, Table 2), but, again, showed only marginally significant local heritability for ER+ BC risk (Table 2, p = 3.52E-22). One potential candidate gene within this locus is *FTO,* which is associated with both age at menarche (Kichaev et al., 2019) and ER+ BC risk (Wu et al., 2024). Alternatively, this may also represent some longer-range linkage between the genes *TOX3* (Wu et al., 2024) and *CASC16* (Jia et al., 2024), which are more strongly associated with ER+ BC risk and are harbored in an adjacent locus which also had substantial local heritability (chromosome 16: 52035823-53382571, 1.6% of total genomic ER+ BC risk, p=1.53E-55, Table S10). Finally, the third locus (chromosome 17: 51,826,118-53,599,431) positively covaried with both ER+ BC risk and age at menarche and had significant local heritability for both traits (Table 3), representing 0.3% and 0.2% of the total trait heritability, respectively. This locus harbors *LINC02073* and *CA10*, which have previously been implicated in studies of age of menarche (Elks et al., 2010). No genes in this locus or neighboring loci have been found to associate with ER+ BC risk, but this locus does harbor *MTCO1P40* and *RPS2P48,* which have been associated with postmenopausal breast cancer risk (Harris et al., 2024), of which ER+ BC is the most common subtype (Yasui & Potter, 1999).

**Figure 3:**
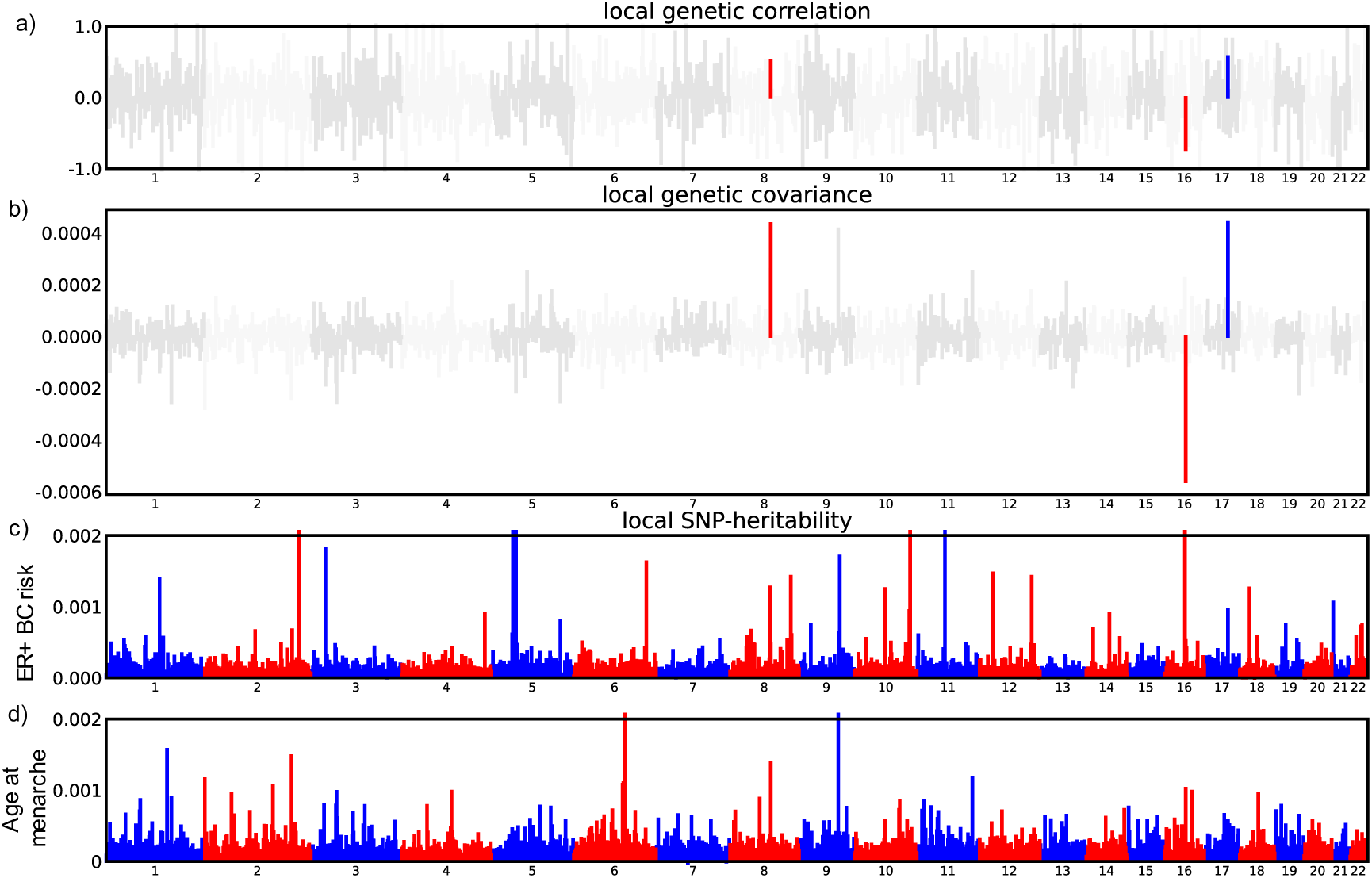
(a) Local genetic correlations, (b) genetic covariance and (c) local heritability for ER+ BC risk and (d) age at menarche across 1,703 loci estimated from population effect GWAS summary statistics. Chromosomes are colored alternatively blue and red. For local genetic covariance and correlations, loci not reaching statistical significance are colored grey. Statistical significance was determined through z-scores after correcting for multiple testing (p < 0.05/1,703).

**Table 2:** Loci with statistically significant local genetic covariance, Local *Cov*(*G_t_*_1,*t*2_), (with standard errors) for ER+ BC risk and age at menarche. Local heritability estimates, Local ℎ^2^ (and standard errors), are bold when reaching statistical significance. All p-values were derived from z-scores, and statistical significance was corrected for multiple testing according to Bonferroni correction and the number of regions tested: p-values < 0.05/1,703 (2.94E-05) were deemed statistically significant. Figure 3 shows these results graphically.

|  | Chromosome 8:76,456,542-79,132,860<br>(n = 4,439 SNPs) |  | Chromosome 16:53,382,572-55,903,773 (n =<br>4,860 SNPs) |  | Chromosome 17: 51826118-53599431<br>(n = 2,958 SNPs) |  |
| --- | --- | --- | --- | --- | --- | --- |
| *Traits* | ER+ BC risk | Age at menarche | ER+ BC risk | Age at menarche | ER+ BC risk | Age at menarche |
| Local $h^2$ | 4.87E-04 (1.53E-04) | **1.39E-03 (1.45E-04)** | 5.24E-04 (1.55E-04) | **1.03E-03 (1.31E-04)** | **9.58E-04 (1.76E-04)** | **5.78E-04 (1.12E-04)** |
| P-value | 7.16E-04 | **5.08E-22** | 3.52E-04 | **2.55E-15** | **2.52E-08** | **1.29E-07** |
| Local $Cov(G_{t1,t2})$ | **4.23E-04 (9.52E-05)** | | **-5.44E-04 (8.78E-05)** | | **4.27E-04 (8.39E-05)** | |
| P-value | **8.59E-06** |  | **5.94E-10** |  | **3.60E-07** |  |

When estimating local heritability and genetic covariance, HESS also estimates genome-wide heritability and genetic correlations. Heritability estimates were in general higher than for LD score regressions (ER+ BC risk = 29.8±0.1%, p < 0.001, *n* = 4,583,461 SNPs; age at menarche = 37.1±0.0%, p < 0.001, *n* = 4,583,461 SNPs; age at first birth = 6.1±0.2%, p < 0.001, *n* = 4,837,687 SNPs), except for number of children (2.8±0.1%, p < 0.001, *n* = 4,862,618 SNPs). Even with the same data, differences in heritability estimates are expected between LD score regressions and HESS because they have slightly different methods of estimating heritability, and HESS does not explicitly account for the effects of population stratification in the GWAS summary stats (Shi et al., 2017). Consequently, while our LD score analyses identified that the inflation of the population effect GWAS data was largely due to polygenicity (Table S4), it is possible that some of our regions of significant local heritability and genetic covariance may not represent direct genetic effects and be false positives due to inflated population effect GWAS data. However, any effects of inflated summary statistics on our core results from HESS analyses appear minimal: our significant local genetic (co)variances coincided with plausible candidate genes, only 1 of the 30 regions with significant local heritability for ER+ BC risk exhibited pleiotropy with any one of the reproductive traits, and only three regions of significant local covariance were detected overall. Finally, even with potentially inflated GWAS data, we repeated the finding of non-statistically significant genome-wide genetic correlations (age at menarche, 0.006±0.008, p = 0.460, *n* = 4,583,461 SNPs, age at first birth, 0.012±0.011, p = 0.270, *n* = 4,837,687 SNPs, number of children = 0.011±0.012, p = 0.372, *n* = 4,862,618 SNPs, Figure 2, yellow, Table S9). Thus, taken together, these analyses suggest no genome-wide genetic correlations between ER+ BC risk and reproductive traits due to largely independent genetic architectures.

## DISCUSSION

We examined the extent and influence of shared genetic architecture between ER+ BC risk and three key reproductive traits (age at menarche, age at first birth, and number of children) to examine evolutionary theories explaining genetic predispositions to a major disease in contemporary European-ancestry populations. Our study builds on research looking at individual loci (Day et al., 2017; Smith et al., 2012; Xiang et al., 2025; Zhao et al., 2024) and estimating genome-wide correlations (Jiang et al., 2019; Prince et al., 2024) by estimating genome-wide correlations while accounting for potential biases in genetic effects (Kong et al., 2018; Young et al., 2022; Young, 2024) and decomposing the genome-wide genetic variation into 1,703 local regions (Berisa & Pickrell, 2016; Shi et al., 2017). While our biobank analyses were underpowered, using the largest GWAS data available, we found no statistically significant genome-wide genetic correlations, regardless of the method used, suggesting that the importance of reproductive traits in shaping genetic ER+ BC risk is low and that common genetic variation does not underlie phenotypic trends observed in epidemiological studies (Aktipis, Ellis, et al., 2015; Kelsey et al., 1993; Momenimovahed & Salehiniya, 2019). Decomposing the genome-wide genetic variation into 1,703 approximately LD-independent genomic regions (Berisa & Pickrell, 2016; Shi et al., 2017) revealed that these results are largely due to a lack of shared genetic architecture between ER+ BC risk and reproductive traits, with the exception of three regions with significant local covariance between age at menarche and ER+ BC risk (Figure 3).

In particular, our analyses suggest that the role of life-history trade-offs in shaping ER+ BC risk is low: only one of the three regions with significant local covariance between reproductive traits and ER+ BC risk was in the direction predicted by life-history theory (i.e., earlier ages at menarche associated with higher ER+ BC risk, chromosome 16: 53,382,572-55,903,773). This suggests that the difference in results between studies focusing only on individual genome-wide significant loci (which show some evidence for life history trade-offs shaping (ER+) BC risk; Day et al., 2017; Smith et al., 2012; Xiang et al., 2025; Zhao et al., 2024) versus studies focusing on whole-genome genetic correlations (which find no evidence, Jiang et al., 2019; Prince et al., 2024) is due to the largely independent genetic architectures of these traits and not due to a symmetrical distribution of significant local genetic covariance (Jiang et al., 2019; van Rheenen et al., 2019). Future studies could further explore GWAS data to identify smaller pleiotropic loci missed in our regional approach and explore which genetic variants causally drive these local covariances using colocalization analysis (Giambartolomei et al., 2014). However, our analyses, which capture all regions of significant local heritability, suggest that even if these variants exist, they will play a small role in shaping genetic predispositions to the disease. Our study focused exclusively on reproductive traits, but body size – due to its impact on an organism’s number of cells – represents another important dimension of evolutionary constraints affecting cancer susceptibility (Aktipis et al., 2013; Nunney, 2018, 2024; Seluanov et al., 2018; Tollis et al., 2020; Vazquez et al., 2022). Height has been identified as a potential risk factor for breast cancer (Green et al., 2011) and – although current evidence of this being due to genetic factors is lacking (Jiang et al., 2019) – it’s role in shaping genetic predispositions to ER+ BC warrants further study.

Our results also provide no evidence for strong, directional selection across ER+ BC risk loci owing to the lack of global and local genetic correlations with ages at first birth and number of children, which are closely related to fitness (Mathieson et al., 2023; Tropf et al., 2015). Thus, while previous phenotypic associations would suggest that ER+ BC risk variants are under negative selection, our results suggest that these phenotypic trends do not translate to the genetic level (Haldane, 1954; Lande & Arnold, 1983; Robertson, 1966). Importantly, while the statistically significant genome-wide heritability for age at first birth and number of children reiterates that contemporary European populations retain the ability to evolve through natural selection (Byars et al., 2010; Fisher, 1930; Haldane, 1954), the heritability was low overall and few regions of significant local heritability were detected (two and zero loci, respectively). As with age at menarche, future studies could explore the role of individual genetic variants with these traits (Giambartolomei et al., 2014), but few loci of large effect have been detected (Barban et al., 2016; Mathieson et al., 2023; Mills et al., 2021) and our approach using HESS (Shi et al., 2017) should be well placed to detect any regions of strong covariance between these traits and ER+ BC risk. It is possible that future methods and larger datasets are better able to detect pleiotropic loci, but if current trends continue, methodological innovations are more likely to remove artificial inflation of heritability estimates rather than discover overlooked genetic variance (Howe et al., 2022; Tropf et al., 2015, this paper, Table 1). Thus, current research suggests that there is little statistical power to detect directional selection on traits by searching for genetic covariation with numbers of children and ages at first birth among common variants in contemporary European populations, although recent work studying rare variants appears more promising (Souaiaia et al., 2026).

Therefore, it remains unclear which evolutionary processes gave rise to genetic predispositions to ER+ BC. Beyond reproductive traits, pleiotropy between diseases acting in early- versus late-life may also maintain genetic variation in disease risk, also in line with the antagonistic pleiotropy theory of ageing (Rodríguez et al., 2017). One potential candidate for ER+ BC risk is Alzheimer’s disease, with higher cancer risk being protective against Alzheimer’s (Ospina-Romero et al., 2020), and recent work illuminating potential underlying mechanisms (Li et al., 2026), which may include genetic factors (Debnath et al., 2026; Ren et al., 2024). However, ER+ BC also has a relatively late age at onset (Yasui & Potter, 1999), suggesting that genetic ER+ BC risk variants may also have accumulated in populations due to weak selection against mutations manifesting at older ages, in line with the mutation accumulation theory of ageing (Charlesworth, 1980; Haldane, 1941; Medawar, 1952). Thus, while our results suggest a minor role for reproductive life-history trade-offs in maintaining genetic variation in ER+ BC risk, studies exploring pleiotropy between ER+ BC and other diseases could help resolve the relative importance of antagonistic pleiotropy and mutation accumulation in shaping genetic variation in this disease.

An important challenge highlighted in this study is the detection of genetic life-history trade-offs in the presence of traits influenced by evolutionary mismatches, as exactly how evolutionary mismatches have influenced both phenotypic and genetic associations between ER+ BC risk and reproductive traits remains unclear. Firstly, while the low heritability of these traits and phenotypic evidence suggest that the phenotypic associations predicted by the mismatch theory are largely acting through environmental factors (e.g., changing cultural preferences), this does not rule out evolutionary mismatches influencing genetic associations through gene-by-environment interactions (Lea et al., 2023). The recent environmental changes in contemporary European populations (Corbett et al., 2018) mean that genetic associations measured in these populations may not represent associations over evolutionary timescales. While the only study – as far as we are aware – looking at gene-by-environment interactions with ER+ BC suggested little overall importance of these effects, the analyses were constrained to the variation in environments experienced within the contemporary European population studied and, even still, detected some loci associated with both ER+ BC risk and reproductive traits which exhibited gene-by-environment interactions (Middha et al., 2023). Thus, studies are needed which quantify the effects of gene-by-environment interactions on reproductive traits and disease beyond the scope of environments experienced by contemporary Europeans.

Conducting longitudinal genomic studies on populations undergoing fertility transitions and other lifestyle changes provides an opportunity to measure the role played by these gene-by-environment interactions and evolutionary mismatches in driving genetic risks for diseases such as cancer (Lea et al., 2023). Moreover, these studies may help unravel the causal mechanisms causing the phenotypic associations between reproductive traits and ER+ BC risk. To the best of our knowledge, the strongest evidence for the role of the evolutionary mismatch remains indirect: for example, reproductive traits are stronger risk factors for ER+ than for estrogen receptor-negative breast cancer (ER- BC, Aktipis, Ellis, et al., 2015) and differences in birth rates among countries predict breast cancer prevalence (You et al., 2018). Moreover, pleiotropic genes involving fertility are likely more easily detected in populations with higher birth rates (e.g., Trumble et al., 2023; and, for breast cancer, Smith et al., 2012). Thus, longitudinal studies on populations undergoing these shifts in reproductive behavior would enable us to gain an understanding of both the role of evolutionary mismatches in shaping genomic associations for important life-history traits and disease, while unravelling potential causal mechanisms of why reproductive traits shape ER+ BC risk. Unfortunately, this research is currently limited by the lack of human diversity reflected in GWAS studies (Mills & Rahal, 2020) and research on understudied populations is needed.

Our study suffers from a few key limitations. First, although we found little evidence in support of life-history trade-offs for ER+ BC risk, exploring genetic correlations between estrogen receptor-negative breast cancer (ER- BC) risk and life-history traits may find different results. While ER- BC risk is not as strongly linked to reproductive life-history traits as ER+ BC risk at the phenotypic level (Aktipis, Ellis, et al., 2015), ER- BC risk does show higher heritability than ER+ BC (Bauer et al., 2007), and there is some evidence that ER- BC risk is genetically correlated with earlier reproduction (Prince et al., 2024), i.e., a fast life-history, as previously hypothesized (Hidaka & Boddy, 2016). Second, we could not account for age at menopause in our analysis due to insufficient sample sizes. Age at menopause is an important life-history trait, and late ages at menopause are a risk factor for ER+ breast cancer (Kelsey & Bernstein, 1996), with previous studies showing some evidence for genetic variants underlying age at menopause also influencing cancer risk (Benonisdottir et al., 2024). Finally, despite using both individual-level biobank data and the largest available GWAS data for life-history traits and ER+ BC risk, some of our genetic correlations were still somewhat imprecise. These power limitations were mostly constrained to the GREML analyses of Lifelines biobank data, where some traits in some analyses did not show significant heritability (e.g., ER+ BC risk and number of children when using the Global screening array and CytoSNP array data). However, even when some of these traits showed higher and statistically significant heritability (e.g., using the Affymetrix data), the finding of non-significant genetic correlations was replicated, although we could only rule out genetic correlations larger than ±0.29-0.52 for the Global screening array data. Estimating genetic correlations with GREML is computationally demanding but further studies could conduct similar analyses using larger biobanks. Nevertheless, our analyses of GWAS data were less power-limited, more precise, and likely less biased. On the whole, our study therefore provides a robust assessment of the strength of genetic correlations and extent of shared genetic architecture between ER+ BC risk and reproductive traits, given current GWAS data and methods available.

In summary, our study uses a multifaceted approach to provide a better understanding of the genetic and evolutionary processes shaping ER+ BC risk in European ancestry populations, which is a deadly form of cancer with increasing prevalence. We encourage future studies to also validate findings across different datasets and approaches using the wealth of methods and genomic data available, but it is important to also broaden the diversity of human genomics and conduct longitudinal studies if we are to unravel the evolutionary process and the underlying mechanisms shaping ER+ BC risk. In particular, our study highlights the challenge of quantifying the influence of evolutionary mismatches on genomics data in contemporary European populations, and assessing the influence of gene-by-environment interactions remains an important line of future research.

## METHODS

### Lifelines biobank

Lifelines is a multi-disciplinary prospective population-based cohort study examining, in a unique three-generation design, the health and health-related behaviors of 167,729 persons living in the North of the Netherlands. It employs a broad range of investigative procedures in assessing the biomedical, socio-demographic, behavioral, physical and psychological factors which contribute to the health and disease of the general population, with a special focus on multi-morbidity and complex genetics. We focused on data from 97,131 women born 1919-1979 and recruited 2007-2013 (https://data-catalogue.lifelines.nl/, accessed 08-05-2024, *Lifelines*, 2024).

### Life-history traits

Age at menarche describes the age in which a female first started menstruating (Rees, 1993), can vary among individuals due to both environmental (e.g., nutritional status) and genetic effects, and is linked to future disease risk, including breast cancer (Parent et al., 2003; Perry et al., 2015). We quantified age at menarche using the lifelines baseline question: “How old were you when you had your first menstruation?” (N=85,210 women). For quality control, we removed 153 values >3 standard deviations from the mean. This gave a mean age at menarche of 13.1 (SD = 1.7, range = 8-18, N = 85,057, Figure S4A), representative of the Netherlands (Parent et al., 2003).

Age at first birth and number of children born are likewise influenced by both genetic factors and environmental factors which are then further constrained by the social environment, cultural background, and any remaining individual preferences (Barban et al., 2016; Mills et al., 2021). We quantified age at first birth and the number of children ever born using questions for a participant’s birth date and the years of birth of her children focusing on full-term pregnancies. For these variables, we used data from baseline questionnaires, supplemented with follow-up questions during wave two of the Lifelines data collection (2014-2018). For age at first birth, we screened the questionnaire data removing any years of birth of children before an individual was born and after an individual had answered the questionnaire (n = 16). We used the birth dates of participants to calculate ages at first birth (mean = 26.9, 95% CIs = 18.9-36.3, Figure S4B) which occurred 1942-2017 (N = 64,779). The number of children ever born was then calculated only for women who were asked for the years of birth of their children after age 45 to avoid biasing the number of childless women and the total number of children born. This gave a mean number of children born of 2.0 (range 0-8, N = 51,672, Figure S4C). All data were handled in R 4.2.2 (2022-10-31, R Core Team, 2020) using *dplyr* 1.1.4 (Wickham et al., 2019).

### ER+ BC diagnoses

ER+ BC diagnoses were determined by linking Lifelines participants to the Dutch Nationwide Pathology Databank (Palga) which contains the national records of histo- and cytopathology, with complete coverage of the Netherlands since 1991 (www.palga.nl, Casparie et al., 2007). The Palga databank details age and dates of diagnoses, and pathology reports describing the subtype, nature, and grade of the tumor (Casparie et al., 2007). Linkage of Lifelines participants to the Palga databank was successfully done for all, except 244 Lifelines participants, producing 10,114 unique pathology reports on breast tumors across 4,310 individuals.

We subtyped these records according to estrogen receptor status (ER-status) using the report conclusions written by pathologists contained in the Palga excerpts (see Supplementary note 1 for further details). According to national Dutch guidelines, ER-status is considered positive when >= 10% of tumor cells are ER+ (IKNL, 2012). We focused on women whose first cancer diagnosis was breast cancer to remove possible metastatic tumors of different origin (n = 628). We also excluded 15 male diagnoses. This left 3,667 Lifelines participants with their first-cancer diagnosis being breast cancer, diagnosed from 1981-2023, and 2,177 ER+ diagnoses (Figure S5), 415 ER- diagnoses, with 1,090 being unable to be assigned an ER status. The ER+ BC diagnoses had a median age of diagnosis of 51 (range = 24-93),

### Genotype data

Genotype data on European-Ancestry Lifelines participants were available on three different platforms: the Illumina CytoSNP-12v2 array (hence CytoSNP array data), the Infinium Global Screening Array MultiEthnic Disease Version 1.0 (hence Global screening array data), and the Affymetrix FinnGen Thermo Fisher Axiom Custom array (hence Affymetrix array data). Full details about the quality control of each genomic dataset and imputation can be found online at: https://wiki.lifelines.nl/. In brief, single nucleotide polymorphisms (SNPs) were filtered according to minimum allele frequency, Hardy-Weinberg equilibrium tests, Plink call rates, and ancestry outliers. Imputation was also performed using Minimac 2012.10.3 (Howie et al., 2012) with *The Genome of The Netherlands* release 5 (The Genome of the Netherlands Consortium, 2014) and the 1000 Genomes phase1 v3 reference panels (The 1000 Genomes Project Consortium, 2010).

We aimed to combine the different Lifelines genotypes in one GREML analysis and therefore harmonized these three datasets using Genotype Harmonizer 1.4.25 (available at https://github.com/molgenis/systemsgenetics/wiki/Genotype-Harmonizer). This software uses linkage disequilibrium patterns to address strand issues, aligning genotype data to one reference dataset, for which we used the Global screening array. The harmonized data contained 1,032,001 variants across 80,045 individuals.

### GREML analyses of Lifelines data

We performed GREML analyses (using GCTA/1.93.2beta-GCCcore-7.3.0, Yang et al., 2011) which leverages the genomic relatedness between distantly related individuals in a population to examine how genetic variation associated with one trait covaries with another (Yang et al., 2010). We first performed a univariate GREML on ER+ BC risk, age at menarche, age at first birth, and number children of ever born to estimate the heritability (ℎ^2^) of each trait, which is the proportion of total phenotypic variance in a trait 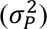 due to genetic factors 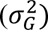:

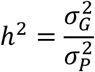

With GREML, the heritability of a given phenotype (**y**) is estimated using a linear mixed model conditioned upon a vector of fixed effects (*_β_*) (such as age and sex) and where the variation explained by the genomic relatedness matrix (*_g_*) is divided by the residual variance (*_ε_*) to estimate the heritability in the equation:

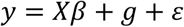

with parameters estimated using the restricted maximum likelihood approach (see Yang et al., 2010, for comprehensive methods).

We then estimated the genetic correlation (*r*(*G*)) between ER+ BC risk and each life-history trait in three bivariate models (Lee et al., 2012). *r*(*G*) is estimated as the genetic covariance between traits *Cov*(*G_t_*_1,*t*2_) scaled by the genetic variance in trait:

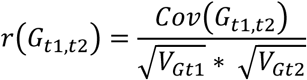

We created the genomic relatedness matrices (GRMs) for all individuals for which we had genomic data using the *--make-grm* argument in the *gcta64* function. We created GRMs for each chromosome first, then merged them into one GRM using --*mgrm*. Finally, the merged GRM was pruned of combinations of closely related individuals while maximizing the sample size using *--grm-cutoff* at a relatedness threshold of 0.05 (*n*-women = 25,068) to remove bias in genetic variances due to gene-environment correlations.

The age of the individual when they were either first diagnosed with breast cancer using the Palga databank, or (for individuals not diagnosed with ER+ BC) when they were last known to be alive and not have ER+ BC, was included as a linear and squared covariate to control for potential age effects while maximizing available sample sizes. The undiagnosed age could be the age of the individual at the date the Palga records were up to date until (February 21^st^, 2023) or the date of death for an individual if this occurred prior (n = 2,293). Dates of death were included in Lifelines as recorded by the Dutch government and were known even in cases of emigration. These dates were decimalized using *lubridate* 1.9.3 in R (Grolemund & Wickham, 2011) and ages were calculated using the birth year and month available in Lifelines. We also included the participant’s birthdate as a linear and squared covariate, to account for any temporal trends in variables, and the top 20 principal components of the pruned GRM used in each model to account for possible population stratification (Yang et al., 2014). Principal components were generated using the --*pca* argument in the *gcta64* function.

However, our bivariate models struggled to converge and univariate models provided apparently biased heritability estimates inconsistent with the literature (e.g., 0% for age at menarche, Table S1d). We posited that this was due to poor coverage of loci between the different genomic samples. For example, although we had over 1 million SNPs in total, the number of SNPs measured in individuals belonging to, for example, both the Affymetrix and CytoSNP array samples was ∼2% of the total number of SNPs available (Table S2). This could result in biased genomic relatedness estimates and increase the chance of missing causal loci, downwardly biasing heritability estimates in the univariate models (Yang et al., 2010). We therefore created GRMs based on each genomic dataset separately where the number of loci measured between individuals was larger and more consistent (Table S2). We present the analysis using the Global screening array data in the main text – because it was the largest in both number of SNPs and individuals – and use the analyses of the other cohorts as replicates. Our sample sizes of informative individuals for our analyses using the Global screening array were: age at menarche, 8,071; age at first birth, 6,260; number of children ever born, 4,602; and 182 ER+ BC cases and 8,700 controls. We also performed a power analysis for our genetic correlations estimated from GREML analyses for each of our traits and genomic datasets based upon heritability estimates from univariate analyses and case and control sample sizes (Figure S1, following Visscher et al., 2014)

Likelihood ratio tests (LRTs) were used to test if heritabilities (in univariate models) and genetic correlations (in bivariate models) were statistically different from 0. For genetic correlations, we adjusted for multiple testing across our three life-history traits using Bonferroni correction to a cutoff of 0.05, giving an adjusted significance threshold of 0.016. To aid convergence, we also used the Expectation–maximization algorithm, increased the number of iterations to 10,000, and standardized life-history trait values to a mean of zero and standard deviation of one.

### GWAS data

We then used LD score regression to also estimate heritabilities and genetic correlations between our traits from previously published GWAS summary statistics (Bulik-Sullivan, Loh, et al., 2015). LD score regression combines genome-wide association study (GWAS) data (i.e., statistical associations between all genetic variants and a trait from a particular study) across different studies to examine how strongly correlated the effects of genetic variants are across different traits, controlling for LD patterns.

For ER+ BC risk, we used GWAS summary statistics from Michailidou et al., (2017), which contained associations of 11,792,542 SNPs with ER+ BC risk based upon 69,501 cases and 105,974 controls. In brief, this study conducted a meta-analytic GWAS by performing a logistic regression on cases and controls using population-based cohort studies, hospital-based studies, and prospective cohorts from contemporary populations, focusing on individuals of European ancestry and controlling for study and genetic ancestry principal components as covariates (see Michailidou et al., 2017, for full methods). We cleaned these data using the *munge_sumstats* function from the software using *ldsc* 1.0.1 (Bulik-Sullivan, Loh, et al., 2015) supported by *Miniconda3*. Specifically, we removed SNPs not found in the HapMap3 to control for imputation quality in the absence of imputation quality scores (*n* = 10,643,464), with MAF < 0.01 (*n* = 10,735), missing values (*n* = 8,108), that were strand ambiguous (*n* = 379), and duplicated identifiers (*n* = 408). This left 1,129,448 SNPs with a mean *_χ_*^2^ value of 1.537, 2,217 of which held genome-wide significance.

For reproductive traits, we first turned to the largest available GWAS data for European ancestry individuals. For age at menarche, we used the data across 11,984,352 SNPs from Loh et al., (2018) conducted on 279,470 European ancestry women in the UK Biobank, and which used a Bayesian regression, including, as covariates, assessment center, genotyping array, age, age squared, and ancestry principal components. For age at first birth, we used meta-analyzed GWAS data across 36 studies totaling 9,744,772 SNPs from Mills et al. (2021) conducted on 542,901 European ancestry individuals and which used linear regressions and included sex, birth year (with polynomials and interactions), top principle components for ancestry, and further cohort-specific covariates, if appropriate, as covariates. Finally, for number of children ever born we used the meta-analyzed GWAS data across 45 cohorts from Mathieson et al., totaling 15,360,087 SNPs (2023) across 785,604 European ancestry individuals, and which also used linear regressions and included, as covariates, birth year (with polynomials), sex, top principal components, further cohort-specific covariates, if appropriate, and for cohorts with family-based data, either controlled for family structure or excluded relatives.

For our reproductive traits, we additionally used GWAS summary statistics from Tan et al. (2024): a recent family-based GWAS study that aimed to separate direct genetic effects of a given SNP on a trait from indirect genetic effects and factors such as assortative mating and population stratification (Kong et al., 2018; Young et al., 2022). This can be achieved by conditioning genetic associations found across individuals upon the parental genotypes in the GWAS model regression; see Tan et al., (2024) for a full description of methods. Tan et al., (2024) estimated the direct genetic effects in a meta-analytic approach across 16 European ancestry cohorts and also accounted for as covariates sex, age (with polynomials and interactions), their interactions, and the first 20 genetic principle components in addition to parental genotypes. The number of SNPs, effective sample sizes, and cohorts studied varied across reproductive traits. Specifically, these were: for age at menarche, *n* = 5,671,230 SNPs, *n* = 19,678 individuals, and four cohorts; for age at first birth: *n* = 6,551,047 SNPs, *n* =35,982, and six cohorts; and for number of children ever born, *n* = 6,412,325 SNPs, *n* = 41,589, and seven cohorts.

For the GWAS data from all reproductive traits, we again used the *munge_sumstats* function to clean the data prior to analyses and, again, selected only sites used in HapMap3, and, for direct genetic effect GWAS data, additionally filtered for SNPs with an effective sample size greater than 80% of the median for each trait to ensure summary statistics were included for only loci whose genetic effects were well estimated (as advised by Tan et al., 2024). For population effect GWAS data, the number of SNPs with known associations for each trait after cleaning were 1,207,271 for age at menarche, 1,194,450 for age at first birth, and 1,202,535 for number of children ever born. The mean and number of genome-wide significant SNPs for these traits were 2.2 and 6,109, 1.6 and 752, and 1.4 and 393, respectively (Table S4). For direct genetic effect GWAS data, the number of SNPs with known associations for each trait after cleaning were 580,219 for age at menarche, 708,027 for age at first birth, and 670,251 for number of children ever born. The mean and number of genome-wide significant SNPs for these traits were 1.127 and 26, 1.078 and 0, and 1.051 and 0, respectively.

### LD score regressions

Heritabilities and genetic correlations were estimated using the software *ldsc* (1.0.1, Bulik-Sullivan, Finucane, et al., 2015; Bulik-Sullivan, Loh, et al., 2015). *ldsc* estimates the heritability of a trait by regressing the *_χ_*^2^-statistics from SNPs on linkage disequilibrium scores using the equation:

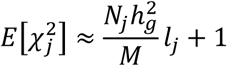

where 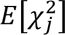 is the expected *_χ_*^2^-statistics for the association between the trait and SNP *j*, *N_j_* is the number of individuals measured for the trait for SNP 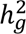 is the squared correlation between the phenotype and the best linear unbiased predictor estimated from the SNPs, *M* is the total number of SNPs, and *l_j_* denotes the LD score of SNP *j* (Bulik-Sullivan, Loh, et al., 2015). In all models, we used global European ancestry LD scores from the 1000 Genome project (The 1000 Genomes Project Consortium, 2010) available from online at: https://zenodo.org/records/7796478. We removed loci in the major histocompatibility complex because of its unusual genetic architecture.

Genome-wide genetic correlations were estimated using the following equation where the expected value of *z*_1*j*_*z_2_l_j_* for a SNP *j* is

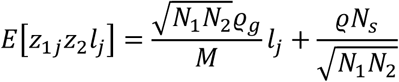

where *N_ig_* is the sample size for the study *i*, *ρ_g_* is the genetic covariance, *_lj_* is the LD score, *_Ns_* is the number of total individuals in both studies, *ρ* is the phenotypic correlation among the overlapping samples (Bulik-Sullivan, Finucane, et al., 2015). We can then normalize the genetic covariance using the heritabilities from trait 1, 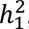, and trait 2, 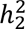, to estimate the genetic correlation, *r_g_*, with the equation:

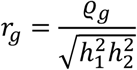

For reproductive traits we used both types of GWAS data outlined above. Significance of the genetic correlations was determined by z-score derived p-values provided by *ldsc,* adjusted to a significance threshold of 0.016 for multiple testing.

### HESS analyses

We used *ρ*-HESS (v0.5, released 9 October 2017, Shi et al., 2017) to detect pleiotropic loci hidden from the genome-wide approaches used by GREML and LD score regressions. For this, we used the previously described population effect GWAS summary statistics (Loh et al., 2018; Mathieson et al., 2023; Michailidou et al., 2017; Mills et al., 2021). We removed duplicate SNPs, SNPs missing statistical associations, and non-autosomal loci. As recommended by HESS, we also only used sites available in the Genomes phase1 v3 reference panels (The 1000 Genomes Project Consortium, 2010) using reference SNP IDs and alleles to merge the data (*n* = 8,676,905 SNPs). All SNPs in this reference panel have a minor allele frequency greater than 1%. Prior to analysis, HESS further filtered for ambiguous strand SNPs (e.g., A/T, C/G) and allele mismatches between GWAS and reference panel. The merged SNP counts for each trait were: for ER+ breast cancer risk, 8,646,068, for age at menarche, 7,757,060; for age at first birth, 8,076,181; and for number of children, 8,204,586.

We used the 1,703 LD independent blocks, on average 1.6 Mb in length, based upon European LD patterns to partition our genome-wide heritability and genetic correlations into local regions (Berisa & Pickrell, 2016). Because the cohorts from Michailidou et al., (2017) was not from any of the biobanks used in the GWAS summary statistics for other traits, we assumed no overlap in the individuals in the analyses. As SNP-level sample sizes were not available we used the number of individuals sampled in the analyses as global sample sizes. The SNP counts for each bivariate analysis between ER+ BC risk and reproductive traits were: for age at menarche, 4,583,461; age at first birth, 4,837,687; and number of children, 4,862,618. P-values were approximated from z-scores, and significance was adjusted for multiple testing across the number of sites p < 0.05/1,703 = 2.9 × 10−5. All local heritability and genetic correlation estimates are available in the online data repository.

## DATA AVAILABILITY

Data may be obtained from Lifelines and are not publicly available. More information about how to request Lifelines data and the conditions of use can be found on their website (https://www.lifelines-biobank.com/researchers/working-with-us/step-1-prepare-and-submit-your-application).

## ACKNOWLEDGEMENTS

We thank the Lifelines data management team and especially Sylvia Gerritsma for their support during this project and Mariana Escobar Rodríguez, Grigory Sivolenko, Thijs Janzen, Raphaël Scherrer, Judith Vonk, Harold Snieder, and Ilja Nolte for their advice on the data handling and analysis. Finally, we would also like to thank the University of Exeter’s quantitative genetics club and Luca G. Hahn for conceptual feedback and discussions regarding the results. We thank the Center for Information Technology of the University of Groningen for their support and for providing access to the Hábrók high performance computing cluster.

## ETHICAL APPROVAL INFORMATION

The Lifelines Cohort Study is conducted following the principles of the Declaration of Helsinki and with approval from the Medical Ethics Committee of the University Medical Center Groningen, Groningen (number 2007/152). Written informed consent was given by each Lifelines participant and the research presented here were approved by Lifelines board (number OV17_00378), adhering to the Lifelines User Code of Conduct. All data was pseudo-anonymized before being received and was handled in accordance with the General Data Protection and Regulation act. This research was approved by the scientific commission of Palga (number 2023-126). Further, data were accessed only within the secure Lifelines workspace or cluster environment on the University Medical Center Groningen cluster *Gearshift* (https://docs.gcc.rug.nl/gearshift/cluster/).

