## supplementary materials for "No genome-wide correlations and little shared genetic architecture between reproductive life-history traits and estrogen receptor-positive breast cancer risk"

Table S1: Results from three univariate GREML models using the **(a)** Global screening, **(b)** Affymetrix, and **(c)** CytoSNP array data examining the genetic and environmental variance of ER+ breast cancer risk and three life-history traits: age at menarche, age at first birth, and the number of children ever born. The table presents parameter estimates (SE) for: $\sigma_G^2$, genetic variance, $\sigma_E^2$, environmental variance; $\sigma_P^2$, phenotypic variance; $h^2$ heritability; $\chi_1^2$, chi-squared test statistic from a LRT test, and corresponding p-value. Values are rounded to two significant digits. Sample sizes for models are shown in the header.

**(a)** Global screening array

| *Traits* | *ER+ BC risk* (n = 182 cases and 8,700 controls) | Age at menarche (n = 8,071) | *Age at first birth* (n = 6,260) | *Number of children* (n = 4,602) |
|---|---|---|---|---|
| $\sigma_G^2$ | 0.0013 (0.00056) | 0.42 (0.058) | 0.21 (0.065) | 0.11 (0.081) |
| $\sigma_E^2$ | 0.0094 (0.00056) | 0.55 (0.055) | 0.66 (0.063) | 0.72 (0.079) |
| $\sigma_P^2$ | 0.011 (0.00016) | 0.97 (0.016) | 0.87 (0.016) | 0.82 (0.017) |
| $h^2$ | 0.12 (0.052) | 0.43 (0.058) | 0.24 (0.074) | 0.13 (0.098) |
| $\chi_1^2$ | 0.77 | 57 | 11 | 0.18 |
| p-value | 0.19 | 1.9e-14 | 0.00058 | 0.34 |

**(b)** Affymetrix array

| *Traits* | *ER+ BC risk* (n = 198 cases, 8,972 controls) | Age at menarche (n = 8,550) | *Age at first birth* (n = 6,177) | *Number of children* (n = 4,435) |
|---|---|---|---|---|
| $\sigma_G^2$ | 0.0018 (0.0007) | 0.46 (0.063) | 0.26 (0.078) | 0.16 (0.095) |
| $\sigma_E^2$ | 0.011 (0.0007) | 0.56 (0.061) | 0.67 (0.076) | 0.67 (0.094) |
| $\sigma_P^2$ | 0.012 (0.00018) | 1.0 (0.016) | 0.92 (0.017) | 0.83 (0.018) |
| $h^2$ | 0.14 (0.056) | 0.45 (0.061) | 0.28 (0.083) | 0.20 (0.11) |
| $\chi_1^2$ | 3.0 | 54 | 11 | 1.8 |
| p-value | 0.041 | 1.2e-13 | 0.00054 | 0.088 |

**(c)** CytoSNP array

| *Traits* | *ER+ BC risk* (n = 195 cases, 6,204) | Age at menarche (n = 6,155) | *Age at first birth* (n = 4,778) | *Number of children* (n = 4,592) |
|---|---|---|---|---|
| $\sigma_G^2$ | 0.0017 (0.00081) | 0.17 (0.053) | 0.17 (0.060) | 0.079 (0.075) |
| $\sigma_E^2$ | 0.014 (0.00083) | 0.83 (0.054) | 0.72 (0.060) | 1 (0.077) |

| | | | | |
|---|---|---|---|---|
| $\sigma_P^2$ | 0.016 (0.00028) | 1.0 (0.018) | 0.88 (0.018) | 1.1 (0.023) |
| $h^2$ | 0.11 (0.05) | 0.17 (0.053) | 0.19 (0.067) | 0.074 (0.069) |
| $\chi_1^2$ | 0.85 | 11 | 6.6 | 0.0 |
| p-value | 0.18 | 0.00050 | 0.0050 | 0.5 |

**(d)** All combined

| *Traits* | *ER+ BC risk* (n = 338 cases, 14,404 controls) | Age at menarche (n = 13,834) | *Age at first birth* (n = 10,418) | *Number of children* (n = 8,284) |
|---|---|---|---|---|
| $\sigma_G^2$ | 0.00061 (0.00019) | 1e-06 (0.019) | 0.00068 (0.023) | 0.12 (0.039) |
| $\sigma_E^2$ | 0.012 (0.00023) | 0.98 (0.022) | 0.93 (0.026) | 1.3 (0.043) |
| $\sigma_P^2$ | 0.013 (0.00015) | 0.98 (0.012) | 0.93 (0.013) | 1.4 (0.022) |
| $h^2$ | 0.048 (0.015) | 1e-06 (0.019) | 0.00073 (0.025) | 0.086 (0.027) |
| $\chi_1^2$ | 6.7 | 0.0010 | 0.25 | 6.8 |
| p-value | 0.0048 | 0.49 | 0.31 | 0.0046 |

Tables S2: Number of quality controlled and harmonized SNPs available for each of the Lifelines genomic datasets (diagonal) and the pairwise-overlap in the number of SNPs between each (below diagonal).

| Genomic data | Global screening array | Affymetrix array | CytoSNP array |
|---|---|---|---|
| Global screening array | 535,107 | | |
| Affymetrix array | 107,735 | 441,001 | |
| CytoSNP array | 67,364 | 21,019 | 244,041 |

Table S3: Results from three bivariate GREML models using the (a) Global screening, (b) Affymetrix, and (c) CytoSNP array genomic data examining the genetic and environmental (co)variance between ER+ breast cancer risk and three life-history traits: age at menarche, age at first birth, and the number of children ever born. The table presents parameter estimates (SE) for: $\sigma_G^2$, genetic variance, $\sigma_E^2$, environmental variance; $\sigma_P^2$, phenotypic variance; $h^2$, heritability; $Cov(G_{t1,t2})$, genetic covariance; $Cov(E_{t1,t2})$, environmental or residual covariance; $r(G_{t1,t2})$, phenotypic covariances which were approximated from genetic and environmental covariances and standard errors (using $\sqrt{(SE_{t1}^2 + SE_{t2}^2)}$); and $r(G_{t1,t2})$, genetic correlation between ER+ breast cancer risk and the life-history traits. Values are rounded to two significant digits. Sample sizes for models are shown in the header.

**(a)** Global screening array

| | ER+ BC risk ~ Age at menarche (n = 176 cases and 7,895 controls) | | ER+ BC risk ~ Age at first birth (n = 155 cases and 6,105 controls) | | ER+ BC risk ~ Number of children (n = 146 cases and 4,456 controls) | |
|---|---|---|---|---|---|---|
| Traits | ER+ BC risk | Age at menarche | ER+ BC risk | Age at first birth | ER+ BC risk | Number of children |
| $\sigma_G^2$ | 0.0013 (0.00056) | 0.42 (0.058) | 0.0014 (0.00057) | 0.23 (0.065) | 0.0012 (0.00056) | 0.16 (0.082) |

| | | | | | | |
|---|---|---|---|---|---|---|
| $\sigma_E^2$ | 0.0094 (0.00056) | 0.55 (0.055) | 0.0093 (0.00056) | 0.64 (0.063) | 0.0095 (0.00056) | 0.66 (0.08) |
| $\sigma_P^2$ | 0.011 (0.00016) | 0.97 (0.016) | 0.011 (0.00016) | 0.87 (0.016) | 0.011 (0.00016) | 0.83 (0.018) |
| $h^2$ | 0.12 (0.053) | 0.43 (0.058) | 0.13 (0.053) | 0.26 (0.074) | 0.11 (0.052) | 0.20 (0.098) |
| $Cov(G_{t1,t2})$ | 0.0012 (0.0041) | | 0.0018 (0.0044) | | 0.0014 (0.0049) | |
| $Cov(E_{t1,t2})$ | 0.0011 (0.004) | | 0.0017 (0.0043) | | 0.0015 (0.0049) | |
| $r(P_{t1,t2})$ | 0.0023 (0.0057) | | 0.0035 (0.0062) | | 0.0029 (0.0069) | |
| $r(G_{t1,t2})$ | 0.051 (0.17) | | 0.098 (0.24) | | 0.10 (0.35) | |
| $\chi_1^2$ | 0 | | 0 | | 0 | |
| p-value | 1.0 | | 1.0 | | 1.0 | |

**(b)** Affymetrix array

| | ER+ BC risk ~ Age at menarche (n = 193 cases and 8,358 controls) | | ER+ BC risk ~ Age at first birth (n = 170 cases and 6,008 controls) | | ER+ BC risk ~ Number of children (n = 145 cases and 4,290 controls) | |
|---|---|---|---|---|---|---|
| Traits | ER+ BC risk | Age at menarche | ER+ BC risk | Age at first birth | ER+ BC risk | Number of children |

| | | | | | | |
|---|---|---|---|---|---|---|
| $\sigma_G^2$ | 0.0021 (0.0007) | 0.46 (0.063) | 0.0019 (0.0007) | 0.28 (0.078) | 0.0017 (0.0007) | 0.21 (0.096) |
| $\sigma_E^2$ | 0.01 (0.0007) | 0.56 (0.061) | 0.01 (0.0007) | 0.65 (0.076) | 0.011 (0.0007) | 0.62 (0.094) |
| $\sigma_P^2$ | 0.012 (0.00018) | 1.0 (0.016) | 0.012 (0.00018) | 0.92 (0.017) | 0.012 (0.00018) | 0.83 (0.018) |
| $h^2$ | 0.17 (0.056) | 0.45 (0.061) | 0.16 (0.056) | 0.30 (0.083) | 0.14 (0.056) | 0.25 (0.11) |
| $Cov(G_{t1,t2})$ | 0.0017 (0.0047) | | 0.0021 (0.0052) | | 0.0014 (0.0058) | |
| $Cov(E_{t1,t2})$ | 0.0016 (0.0046) | | 0.002 (0.0052) | | 0.0014 (0.0058) | |
| $r(P_{t1,t2})$ | 0.0033 (0.0066) | | 0.0041 (0.0074) | | 0.0028 (0.0082) | |
| $r(G_{t1,t2})$ | 0.054 (0.15) | | 0.090 (0.23) | | 0.075 (0.31) | |
| $\chi_1^2$ | 0 | | 0 | | 0 | |
| p-value | 1.0 | | 1.0 | | 1.0 | |

**(c)** CytoSNP array

| | ER+ BC risk ~ Age at menarche (n = 190 cases and 5,965 controls) | ER+ BC risk ~ Age at first birth (n = 147 cases and 4,631 controls) | ER+ BC risk ~ Number of children (n = 176 cases and 4,416 controls) |
|---|---|---|---|

| Traits | ER+ BC risk | Age at menarche | ER+ BC risk | Age at first birth | ER+ BC risk | Number of children |
|---|---|---|---|---|---|---|
| $\sigma_G^2$ | 0.0016 (0.00081) | 0.17 (0.053) | 0.0017 (0.00081) | 0.18 (0.06) | 0.0012 (0.00056) | 0.16 (0.082) |
| $\sigma_E^2$ | 0.014 (0.00083) | 0.83 (0.054) | 0.014 (0.00083) | 0.71 (0.061) | 0.0095 (0.00056) | 0.66 (0.08) |
| $\sigma_P^2$ | 0.016 (0.00028) | 1.0 (0.018) | 0.016 (0.00028) | 0.89 (0.018) | 0.011 (0.00016) | 0.83 (0.018) |
| $h^2$ | 0.10 (0.05) | 0.17 (0.053) | 0.11 (0.05) | 0.20 (0.068) | 0.11 (0.052) | 0.20 (0.098) |
| $Cov(G_{t1,t2})$ | 0.0013 (0.0046) | | 0.0025 (0.0049) | | 0.0013 (0.0055) | |
| $Cov(E_{t1,t2})$ | 0.0013 (0.0047) | | 0.0024 (0.0051) | | 0.0013 (0.0057) | |
| $r(P_{t1,t2})$ | 0.0026 (0.0066) | | 0.0049 (0.0073) | | 0.0026 (0.0081) | |
| $r(G_{t1,t2})$ | 0.078 (0.27) | | 0.14 (0.28) | | 0.091 (0.39) | |
| $\chi_1^2$ | 0 | | 0 | | 0 | |
| p-value | 1.0 | | 1.0 | | 1.0 | |

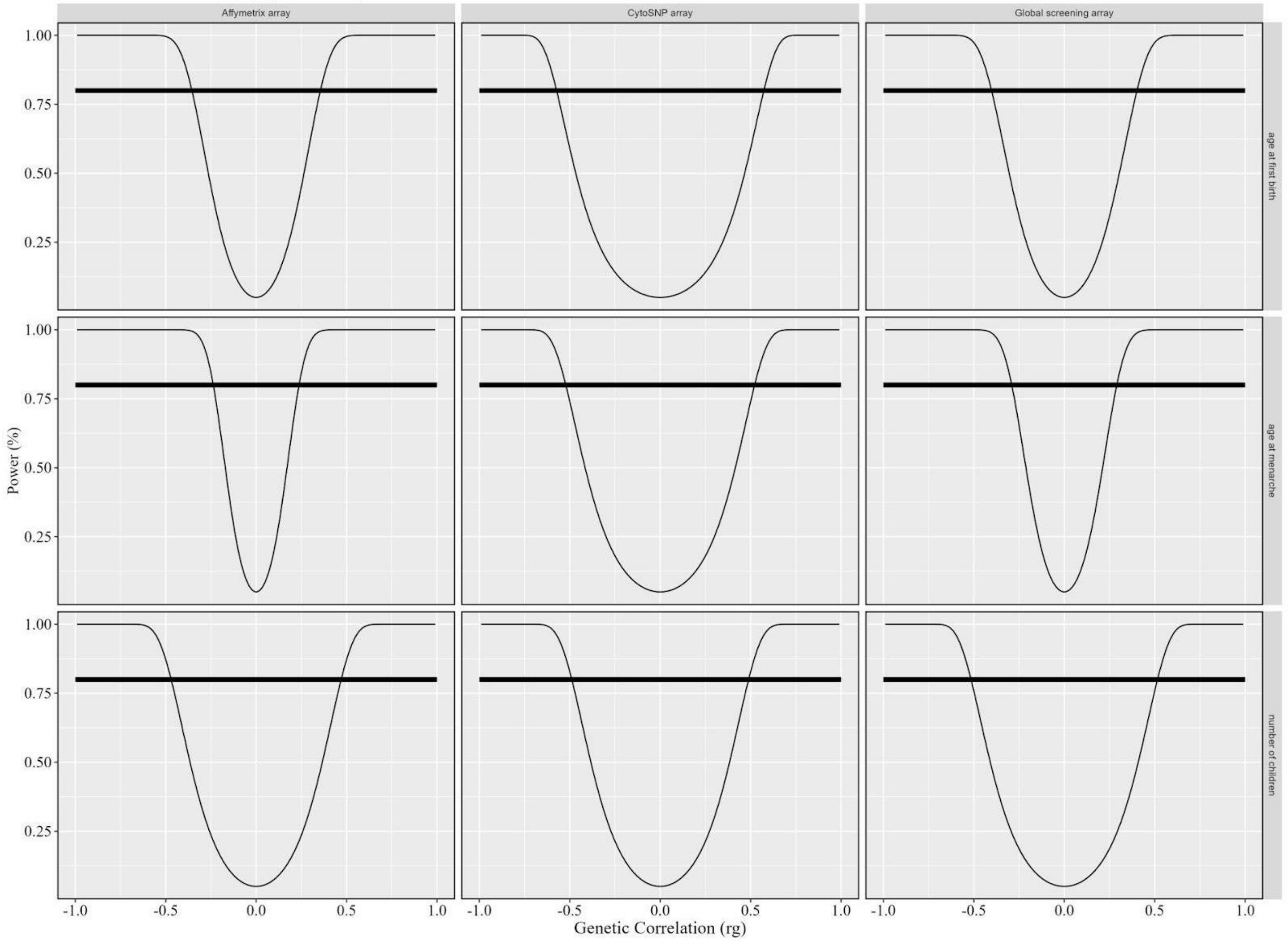

Figure S1: Power to estimate genetic correlations using the Lifelines biobank Affymetrix Global screening array (column 1), CytoSNP array (column 2), and Global screening array data (column 3) between ER+ BC risk and age at first birth (row 1), age at menarche (row 2), and number of children (row 3). Thin line shows the estimated percentage power to estimate genetic correlations ($r_g$) of different sizes from -1 to 1 based on expected population prevalence of 8% and heritability estimates from univariate analyses and case and control sample sizes available in Lifelines for women for each of the genomic datasets. Thick line shows 80% power. When the thin line is above the thick line, genetic correlations of this size could be detected with 80% power.

Table S4: Summary statistics of GWAS data using LD Score Regression with the software *ldsc*. Presented are: the links to datasets, number of SNPs analysed; Mean chi^2 values, Lambda genetic control, Max chi^2, the number of genome-wide significant variants, the heritability estimates (h2) and their standard errors and the *ldsc* intercept and ldsc ratio values, with higher values reflecting potential confounding. Standard errors are shown in parentheses. Values are rounded to three significant digits.

| | Population effects | | | | Direct genetic effects | | |
|---|---|---|---|---|---|---|---|
| Parameter | ER+ BC risk | Age at menarche | Age at first birth | Number of children | Age at menarche | Age at first birth | Number of children |
| Available at: | https://www.ebi.ac.uk/gwas/studies/GCST004988 | https://www.ebi.ac.uk/gwas/studies/GCST90029036 | https://www.ebi.ac.uk/gwas/studies/GCST90000050 | https://www.repository.cam.ac.uk/handle/1810/341056 | All available at: https://thessgac.com/ | | |
| n SNPs | 1129448 | 1207271 | 1194450 | 1202535 | 580219 | 708027 | 670251 |
| Mean chi^2 | 1.537 | 2.2 | 1.615 | 1.449 | 1.127 | 1.078 | 1.051 |
| Lambda GC | 1.276 | 1.694 | 1.467 | 1.353 | 1.111 | 1.066 | 1.057 |
| Max chi^2 | 1424.989 | 197.9 | 143.036 | 103.133 | 93.032 | 23.042 | 20.639 |
| Genome-wide significant SNPs | 2217 | 6109 | 752 | 393 | 26 | 0 | 0 |
| h2 | 0.1004 | 0.1997 | 0.0426 | 0.022 | 0.0929 | 0.0322 | 0.0265 |

| | | | | | | | |
|---|---|---|---|---|---|---|---|
| | (0.0103) | (0.0083) | (0.0019) | (0.0011) | (0.0251) | (0.0098) | (0.0079) |
| Intercept | 1.1261 (0.0125) | 1.1339 (0.0125) | 1.0851 (0.0094) | 1.0575 (0.0082) | 1.0608 (0.0106) | 1.0411 (0.0083) | 1.0148 (0.0074) |
| Ratio | 0.2357 (0.0234) | 0.111 (0.0103) | 0.1385 (0.0153) | 0.1279 (0.0183) | 0.4801 (0.084) | 0.5282 (0.1061) | 0.2959 (0.1473) |

Table S5: Results from three bivariate LD score regressions of population effect GWAS data examining the genetic and environmental (co)variance between ER+ breast cancer risk and three life-history traits: age at menarche, age at first birth, and the number of children ever born. The table presents parameter estimates (SE) for: $h^2$, heritability; $Cov(G_{t1,t2})$, genetic covariance; $mean(Z_{t1 \times t2})$, the average product of GWAS scores across loci between ER+ BC risk and life-history traits; $Cov(G_{t1,t2})$ intercept, genetic covariance intercept from the LD score regression; and $r(G_{t1,t2})$, genetic correlation between ER+ breast cancer risk and the life-history traits, Z score of this correlation, and derived P value. Sample size of SNPs used are shown and other values are rounded to two significant digits.

| | ER+ BC risk ~ Age at menarche (N = 1,116,653 SNPs) | | ER+ BC risk ~ Age at first birth (N = 1,113,353 SNPs) | | ER+ BC risk ~ Number of children (N = 1,114,830 SNPs) | |
|---|---|---|---|---|---|---|
| *Traits* | *ER+ BC risk* | Age at menarche | *ER+ BC risk* | *Age at first birth* | *ER+ BC risk* | *Number of children* |
| $h^2$ | 0.093 (0.011) | 0.213 (0.010) | 0.093 (0.011) | 0.044 (0.002) | 0.093 (0.011) | 0.023 (0.001) |
| $Cov(G_{t1,t2})$ | 0.0000 (0.0034) | | -0.0008 (0.0019) | | -0.0022 (0.0013) | |
| $mean(Z_{t1 \times t2})$ | 0.0047 | | -0.0024 | | -0.0222 | |
| $Cov(G_{t1,t2})$ intercept | -0.0062 (0.0065) | | 0.0019 (0.0058) | | 0.0002 (0.0054) | |
| $r(G_{t1,t2})$ | 0.0003 (0.0239) | | -0.0128 (0.0293) | | -0.0472 (0.0287) | |
| Z score | 0.0126 | | -0.4369 | | -1.6446 | |
| p-value | 0.9900 | | 0.6622 | | 0.1001 | |

Table S6: Results from three bivariate LD score regressions of direct genetic effect GWAS data examining the genetic and environmental (co)variance between ER+ breast cancer risk and three life-history traits: age at menarche, age at first birth, and the number of children ever born. The table presents parameter estimates (SE) for: $h^2$, heritability; $Cov(G_{t1,t2})$, genetic covariance; $mean(Z_{t1 \times t2})$, the average product of GWAS scores across loci between ER+ BC risk and life-history traits; $Cov(G_{t1,t2})$ intercept, genetic covariance intercept from the LD score regression; and $r(G_{t1,t2})$, genetic correlation between ER+ breast cancer risk and the life-history traits, Z score of this correlation, and derived P value. Sample size of SNPs used are shown and other values are rounded to two significant digits. Sample size of SNPs used are shown and other values are rounded to two significant digits.

| | ER+ BC risk ~ Age at menarche (N = 551,453 SNPs) | | ER+ BC risk ~ Age at first birth (N = 668,389 SNPs) | | ER+ BC risk ~ Number of children (n = 635,154 SNPs) | |
|---|---|---|---|---|---|---|
| *Traits* | *ER+risk* | Age at menarche | *ER+ BC risk* | *Age at first birth* | *ER+ BC risk* | *Number of children* |
| $h^2$ | 0.077 (0.015) | 0.108 (0.024) | 0.080 (0.013) | 0.034 (0.010) | 0.079 (0.016) | 0.027 (0.008) |
| $Cov(G_{t1,t2})$ | 0.0027 (0.0085) | | -0.0010 (0.0049) | | -0.0018 (0.0039) | |
| $mean(Z_{t1 \times t2})$ | -0.0082 | | -0.0014 | | -0.0078 | |
| $Cov(G_{t1,t2})$ intercept | -0.0143 (0.0085) | | 0.0014 (0.0061) | | -0.0034 (0.0061) | |
| $r(G_{t1,t2})$ | 0.0291 (0.0944) | | -0.0191 (0.0958) | | -0.0379 (0.0834) | |
| Z score | 0.3083 | | -0.1994 | | -0.4544 | |
| p-value | 0.7579 | | 0.8420 | | 0.6495 | |

Table S7: Results from three bivariate LD score regressions of population effect GWAS data between reproductive traits: age at menarche, age at first birth, and the number of children ever born. The table presents parameter estimates (SE) for: $h^2$, heritability; $Cov(G_{t1,t2})$, genetic covariance; $mean(Z_{t1 \times t2})$, the average product of GWAS scores across loci between ER+ BC risk and life-history traits; $Cov(G_{t1,t2})$ intercept, genetic covariance intercept from the LD score regression; and $r(G_{t1,t2})$, genetic correlation between ER+ breast cancer risk and the life-history traits, Z score of this correlation, and derived P value. Sample size of SNPs used are shown and other values are rounded to two significant digits. Sample size of SNPs used are shown and other values are rounded to two significant digits.

| | age at first birth ~ age at menarche (N = 1,178,825 SNPs) | | age at first birth ~ number of children (N = 1,179,108 SNPs) | | number of children ~ age at menarche (N = 1,183,526 SNPs) | |
|---|---|---|---|---|---|---|
| *Traits* | age at first birth | age at menarche | age at first birth | number of children | number of children | age at menarche |
| $h^2$ | 0.043 (0.002) | 0.210 (0.010) | 0.043 (0.002) | 0.022 (0.001) | 0.022 (0.001) | 0.211 (0.010) |
| $Cov(G_{t1,t2})$ | 0.009 (0.002) | | -0.022 (0.001) | | 0.003 (0.001) | |
| $mean(Z_{t1 \times t2})$ | 0.083 | | -0.484 | | 0.040 | |
| $Cov(G_{t1,t2})$ intercept | 0.023 (0.0063) | | -0.1605 (0.0069) | | 0.0078 (0.0057) | |
| $r(G_{t1,t2})$ | 0.097 (0.021) | | -0.705 (0.021) | | 0.039 (0.020) | |
| Z score | 4.704 | | -33.737 | | 1.911 | |
| p-value | 0.000 | | 0.000 | | 0.056 | |

Table S8: Results from three bivariate LD score regressions of direct genetic effect GWAS data between reproductive traits: age at menarche, age at first birth, and the number of children ever born. The table presents parameter estimates (SE) for: $h^2$, heritability; $Cov(G_{t1,t2})$, genetic covariance; $mean(Z_{t1 \times t2})$, the average product of GWAS scores across loci between ER+ BC risk and life-history traits; $Cov(G_{t1,t2})$ intercept, genetic covariance intercept from the LD score regression; and $r(G_{t1,t2})$, genetic correlation between ER+ breast cancer risk and the life-history traits, Z score of this correlation, and derived P value. Sample size of SNPs used are shown and other values are rounded to two significant digits. Sample size of SNPs used are shown and other values are rounded to two significant digits.

| | age at first birth ~ age at menarche (N = 435,982 SNPs s) | | age at first birth ~ number of children (N = 651,960 SNPs) | | number of children ~ age at menarche (N = 439,322 SNPs) | |
|---|---|---|---|---|---|---|
| *Traits* | age at first birth | age at menarche | age at first birth | number of children | number of children | age at menarche |
| $h^2$ | 0.030 (0.011) | 0.083 (0.024) | 0.030 (0.009) | 0.026 (0.008) | 0.0285 (0.0078) | 0.0821 (0.0237) |
| $Cov(G_{t1,t2})$ | 0.010 (0.010) | | -0.008 (0.006) | | 0.0005 (0.0096) | |
| $mean(Z_{t1 \times t2})$ | 0.030 | | -0.136 | | 0.0038 | |
| $Cov(G_{t1,t2})$ intercept | 0.0204 (0.0065) | | -0.1239 (0.0055) | | 0.0024 (0.0072) | |
| $r(G_{t1,t2})$ | 0.204 (0.216) | | -0.302 (0.210) | | 0.0098 0.1980) | |
| Z score | 0.946 | | -1.438 | | 0.0495 | |
| p-value | 0.344 | | 0.150 | | 0.9605 | |

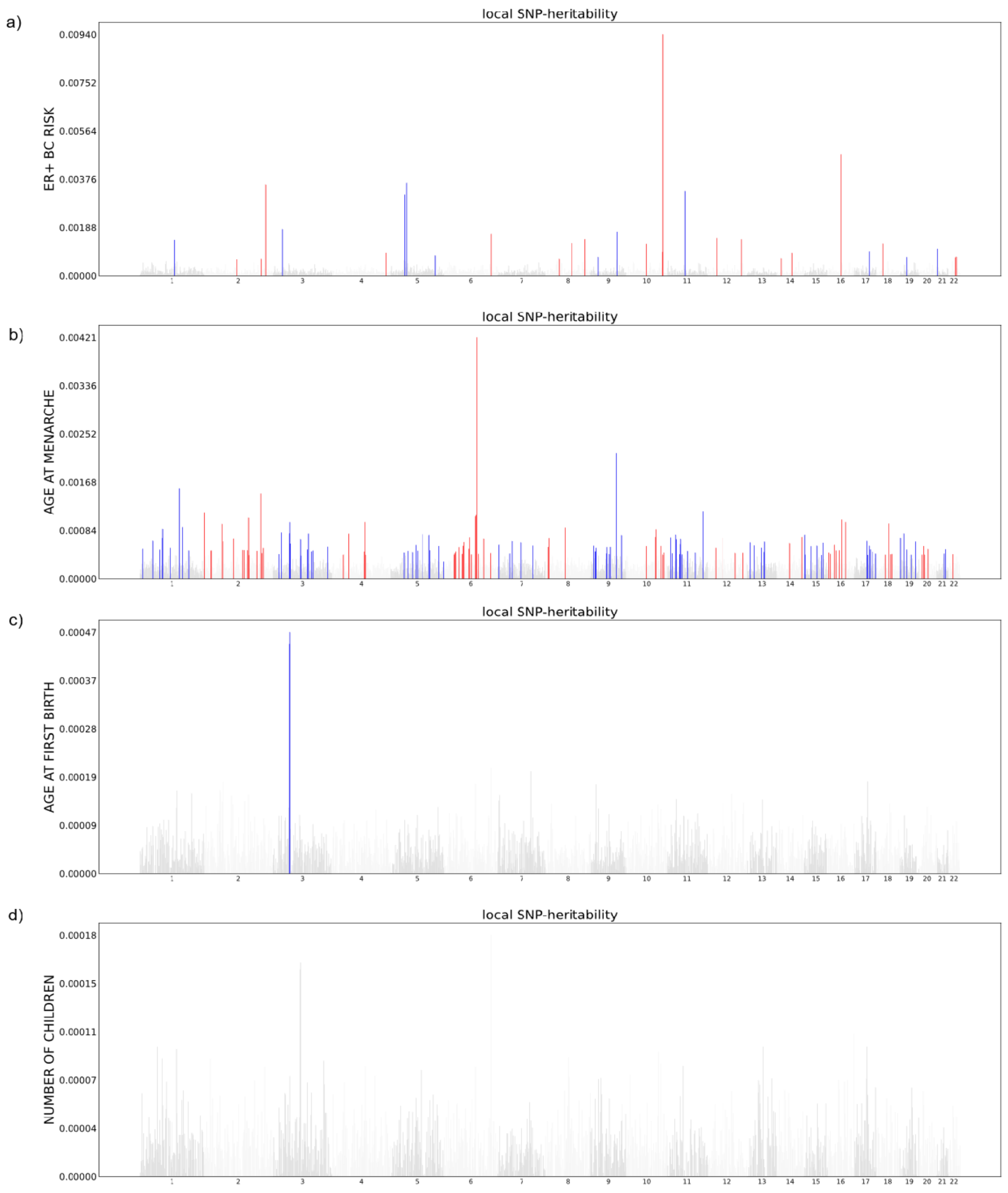

Figure S2: Local heritability across 1,703 loci estimated from population effect GWAS summary statistics for: (a) ER+ BC risk, (b) age at menarche, (c) age at first birth and (d) number of children Chromosomes are colored alternatively blue and red. Loci not reaching statistical significance are colored grey. Statistical significance was determined through z-scores after correcting for multiple testing ($p < 0.05/1{,}703$). Noted that the y axes are scaled according to each trait.

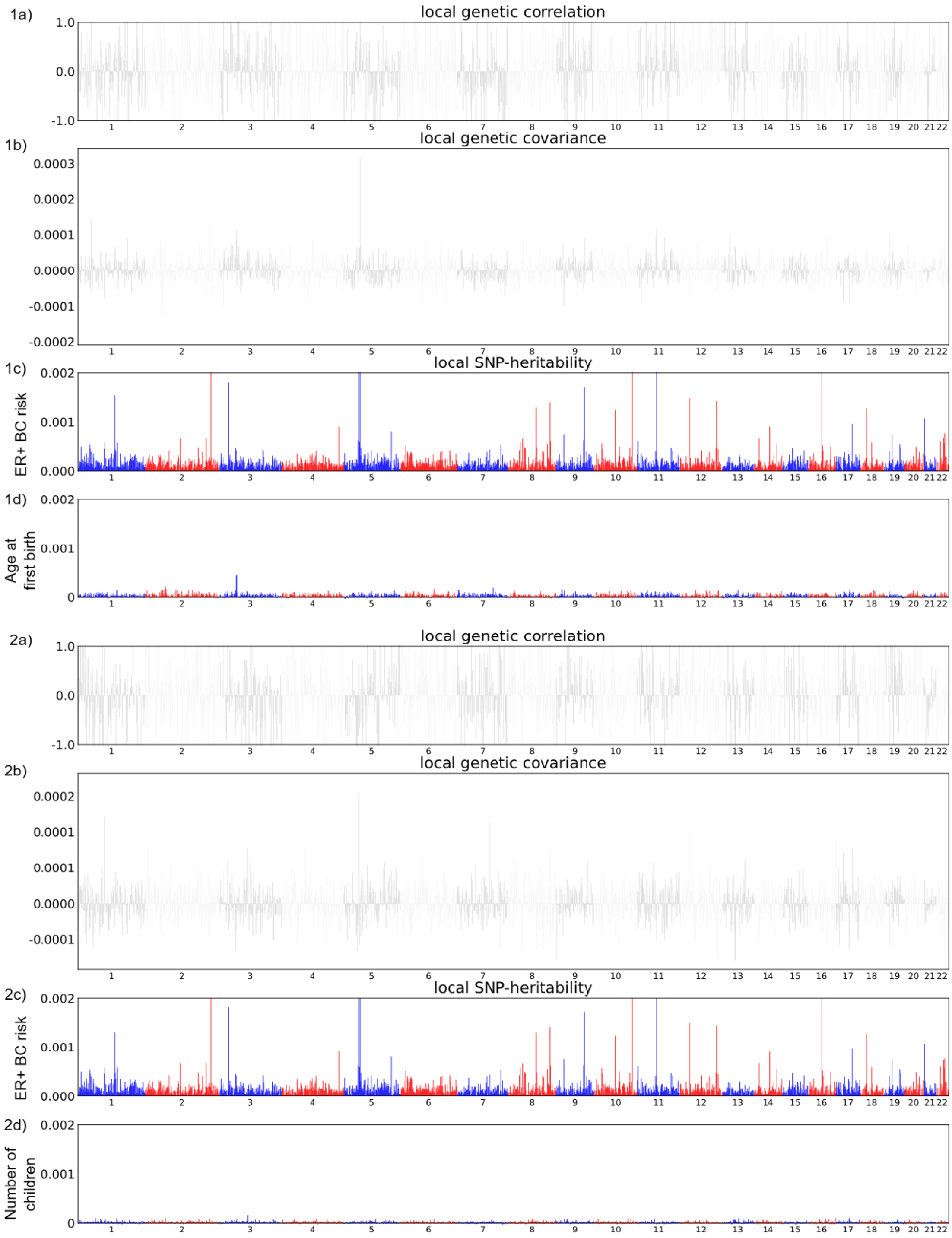

Figure S3: (a) Local genetic correlations, (b) genetic covariance and (c) local heritability for ER+ BC risk and (d) the reproductive traits (1) age at first birth and (2) number of children across 1,703 loci estimated from population effect GWAS summary statistics. Chromosomes are colored alternatively blue and red. For local genetic covariance and correlations, loci not reaching statistical significance are colored grey. Statistical significance was determined through z-scores after correcting for multiple testing ($p < 0.05/1,703$).

Table S9: Results from three bivariate Heritability Estimations from Summary Statistics (HESS) models of GWAS population genetic effects estimating the genome-wide genetic (co)variance between ER+ breast cancer risk and three life-history traits: age at menarche, age at first birth, and the number of children ever born. The table presents parameter estimates (SE) for: $h^2$, heritability; $Cov(G_{t1,t2})$, genetic covariance; and $r(G_{t1,t2})$, genetic correlation between ER+ breast cancer risk and the life-history traits, Z score of this correlation, and derived P value. Sample size of SNPs used are shown and other values are rounded to two significant digits. Sample size of SNPs used are shown and other values are rounded to two significant digits.

| | ER+ BC risk ~ Age at menarche (N = 4,583,461) | | ER+ BC risk ~ Age at first birth (N = 4,837,687) | | ER+ BC risk ~ Number of children (n = 4,862,618) | |
|---|---|---|---|---|---|---|
| *Traits* | *ER+ BC risk* | Age at menarche | *ER+ BC risk* | *Age at first birth* | *ER+ BC risk* | *Number of children* |
| $h^2$ | 0.298 (0.00551) | 0.371 (0.00383) | 0.298 (0.00552) | 0.0605 (0.0023) | 0.296 (0.00553) | 0.0276 (0.00162) |
| $Cov(G_{t1,t2})$ | 0.0019 (0.00244) | | 0.00157 (0.00211) | | 0.000976 (0.0018) | |
| $r(G_{t1,t2})$ | 0.00573 (0.00775) | | 0.0117 (0.0106) | | 0.0108 (0.0121) | |
| Z score | 0.7394 | | 1.1038 | | 0.8926 | |
| p-value | 0.4597 | | 0.2697 | | 0.3721 | |

Table S10: Loci with statistically significant local heritability for ER+ BC risk and reproductive traits. P-values were corrected for multiple testing according to number of regions tested and Bonferroni correction: p values < 0.05/1,703 were deemed statistically significant. All local heritability and genetic correlation estimates are available in the online data repository.

| Chromosome | Loci start position | Loci end position | *n SNPs* | Local $h^2$ | Standard error | Z-score | P-value | Local $h^2$ (%) |
|---|---|---|---|---|---|---|---|---|
| **a) ER+ BC risk** | | | | | | | | |
| 1 | 118839067 | 144977493 | 2894 | 1.40E-03 | 1.95E-04 | 7.180 | 3.50E-13 | 4.70E-03 |
| 2 | 118367466 | 121303782 | 4688 | 6.62E-04 | 1.62E-04 | 4.097 | 2.10E-05 | 2.22E-03 |
| 2 | 201576284 | 202818636 | 1335 | 6.74E-04 | 1.62E-04 | 4.155 | 1.63E-05 | 2.26E-03 |
| 2 | 217715661 | 218395479 | 1182 | 3.55E-03 | 2.69E-04 | 13.173 | 6.30E-40 | 1.19E-02 |
| 3 | 26877769 | 27840909 | 1724 | 1.81E-03 | 2.11E-04 | 8.582 | 4.67E-18 | 6.09E-03 |
| 4 | 174264132 | 176570715 | 3804 | 9.09E-04 | 1.74E-04 | 5.237 | 8.15E-08 | 3.05E-03 |
| 5 | 43983499 | 50163397 | 4069 | 3.17E-03 | 2.58E-04 | 12.286 | 5.40E-35 | 1.06E-02 |
| 5 | 55417349 | 56621101 | 2153 | 3.62E-03 | 2.71E-04 | 13.323 | 8.51E-41 | 1.21E-02 |
| 5 | 156628700 | 158825697 | 4007 | 8.02E-04 | 1.68E-04 | 4.759 | 9.75E-07 | 2.69E-03 |
| 6 | 151912703 | 153094495 | 2279 | 1.63E-03 | 2.04E-04 | 7.983 | 7.16E-16 | 5.47E-03 |
| 8 | 29327896 | 31133728 | 2681 | 6.69E-04 | 1.62E-04 | 4.131 | 1.81E-05 | 2.25E-03 |
| 8 | 75445064 | 76456541 | 1592 | 1.28E-03 | 1.85E-04 | 6.884 | 2.90E-12 | 4.28E-03 |
| 8 | 126410917 | 128659110 | 4336 | 1.43E-03 | 1.96E-04 | 7.279 | 1.68E-13 | 4.79E-03 |
| 9 | 20463534 | 22206558 | 2782 | 7.48E-04 | 1.66E-04 | 4.509 | 3.26E-06 | 2.51E-03 |
| 9 | 110695062 | 112778023 | 4449 | 1.71E-03 | 2.07E-04 | 8.254 | 7.65E-17 | 5.75E-03 |
| 10 | 63341695 | 65794113 | 4108 | 1.25E-03 | 1.89E-04 | 6.639 | 1.58E-11 | 4.21E-03 |
| 10 | 122407323 | 123231464 | 1554 | 9.46E-04 | 1.75E-04 | 5.397 | 3.38E-08 | 3.17E-03 |
| 10 | 123231465 | 123900544 | 1414 | 9.40E-03 | 4.08E-04 | 23.015 | 1.64E-117 | 3.15E-02 |
| 11 | 68005825 | 69516129 | 2436 | 3.31E-03 | 2.62E-04 | 12.617 | 8.51E-37 | 1.11E-02 |
| 12 | 27799773 | 29651254 | 3483 | 1.48E-03 | 1.98E-04 | 7.449 | 4.69E-14 | 4.95E-03 |
| 12 | 115503216 | 117087470 | 2954 | 1.43E-03 | 1.96E-04 | 7.281 | 1.66E-13 | 4.79E-03 |
| 14 | 35859593 | 38667724 | 3665 | 6.97E-04 | 1.63E-04 | 4.263 | 1.01E-05 | 2.34E-03 |
| 14 | 67992317 | 71131956 | 5344 | 9.03E-04 | 1.73E-04 | 5.213 | 9.30E-08 | 3.03E-03 |
| 16 | 52035823 | 53382571 | 2126 | 4.74E-03 | 3.03E-04 | 15.655 | 1.53E-55 | 1.59E-02 |
| 17 | 51826118 | 53599431 | 2958 | 9.58E-04 | 1.76E-04 | 5.450 | 2.52E-08 | 3.22E-03 |
| 18 | 24026191 | 25927681 | 2886 | 1.26E-03 | 1.89E-04 | 6.676 | 1.23E-11 | 4.24E-03 |
| 19 | 18409862 | 19877470 | 1993 | 7.45E-04 | 1.66E-04 | 4.494 | 3.50E-06 | 2.50E-03 |
| 21 | 15950982 | 18053164 | 3062 | 1.07E-03 | 1.81E-04 | 5.899 | 1.83E-09 | 3.58E-03 |
| 22 | 37570269 | 39307893 | 2529 | 7.28E-04 | 1.65E-04 | 4.415 | 5.05E-06 | 2.44E-03 |
| 22 | 40545797 | 42690817 | 2412 | 7.56E-04 | 1.66E-04 | 4.551 | 2.67E-06 | 2.54E-03 |
| **b)** Age at menarche | | | | | | | | |
| 1 | 7247335 | 9365198 | 3463 | 5.27E-04 | 1.10E-04 | 4.801 | 7.88E-07 | 1.42E-03 |
| 1 | 43758457 | 44969182 | 1492 | 6.64E-04 | 1.16E-04 | 5.724 | 5.19E-09 | 1.79E-03 |
| 1 | 65041704 | 66939403 | 2901 | 5.12E-04 | 1.09E-04 | 4.698 | 1.32E-06 | 1.38E-03 |

| | | | | | | | | |
|---|---|---|---|---|---|---|---|---|
| 1 | 71684405 | 74326906 | 3363 | 7.10E-04 | 1.18E-04 | 6.014 | 9.03E-10 | 1.91E-03 |
| 1 | 74326907 | 76728134 | 3786 | 8.68E-04 | 1.25E-04 | 6.948 | 1.85E-12 | 2.34E-03 |
| 1 | 102041016 | 102898744 | 1574 | 5.41E-04 | 1.05E-04 | 5.139 | 1.38E-07 | 1.46E-03 |
| 1 | 165191702 | 166460516 | 2270 | 1.57E-03 | 1.52E-04 | 10.390 | 1.38E-25 | 4.24E-03 |
| 1 | 177433381 | 178944308 | 2172 | 8.97E-04 | 1.26E-04 | 7.112 | 5.71E-13 | 2.42E-03 |
| 1 | 199239884 | 200137648 | 1906 | 4.97E-04 | 1.08E-04 | 4.587 | 2.24E-06 | 1.34E-03 |
| 2 | 10133 | 1781021 | 4416 | 1.16E-03 | 1.37E-04 | 8.493 | 1.01E-17 | 3.12E-03 |
| 2 | 23341383 | 24686917 | 2222 | 4.89E-04 | 1.08E-04 | 4.529 | 2.96E-06 | 1.32E-03 |
| 2 | 24686918 | 26894984 | 2736 | 4.98E-04 | 1.08E-04 | 4.600 | 2.11E-06 | 1.34E-03 |
| 2 | 56203345 | 57429099 | 2087 | 9.53E-04 | 1.28E-04 | 7.423 | 5.75E-14 | 2.57E-03 |
| 2 | 58297315 | 60291999 | 2758 | 5.47E-04 | 1.11E-04 | 4.942 | 3.87E-07 | 1.47E-03 |
| 2 | 60292000 | 62429043 | 2658 | 6.54E-04 | 1.16E-04 | 5.660 | 7.57E-09 | 1.76E-03 |
| 2 | 105125034 | 106210057 | 1926 | 7.02E-04 | 1.18E-04 | 5.959 | 1.27E-09 | 1.89E-03 |
| 2 | 137042794 | 138698116 | 3134 | 5.06E-04 | 1.09E-04 | 4.651 | 1.65E-06 | 1.36E-03 |
| 2 | 141805351 | 142518601 | 1650 | 5.04E-04 | 1.08E-04 | 4.652 | 1.64E-06 | 1.36E-03 |
| 2 | 152486782 | 154005055 | 2372 | 5.01E-04 | 1.08E-04 | 4.617 | 1.95E-06 | 1.35E-03 |
| 2 | 155391135 | 157560634 | 2956 | 1.06E-03 | 1.33E-04 | 7.999 | 6.26E-16 | 2.86E-03 |
| 2 | 157560635 | 158533217 | 708 | 4.13E-04 | 9.35E-05 | 4.420 | 4.94E-06 | 1.11E-03 |
| 2 | 182266031 | 184357607 | 3226 | 4.90E-04 | 1.08E-04 | 4.535 | 2.88E-06 | 1.32E-03 |
| 2 | 199311125 | 201576283 | 2498 | 1.49E-03 | 1.48E-04 | 10.006 | 7.19E-24 | 4.00E-03 |
| 2 | 202818637 | 205799240 | 3546 | 4.52E-04 | 1.06E-04 | 4.260 | 1.02E-05 | 1.22E-03 |
| 2 | 208645399 | 209941528 | 1834 | 5.38E-04 | 1.10E-04 | 4.876 | 5.40E-07 | 1.45E-03 |
| 3 | 17891118 | 19125143 | 1819 | 4.34E-04 | 1.05E-04 | 4.128 | 1.83E-05 | 1.17E-03 |
| 3 | 23804865 | 25461557 | 3371 | 8.03E-04 | 1.22E-04 | 6.574 | 2.45E-11 | 2.16E-03 |
| 3 | 47727212 | 49316971 | 970 | 7.91E-04 | 1.11E-04 | 7.151 | 4.31E-13 | 2.13E-03 |
| 3 | 49316972 | 51832014 | 2008 | 9.83E-04 | 1.28E-04 | 7.691 | 7.29E-15 | 2.65E-03 |
| 3 | 51832015 | 54081389 | 2664 | 6.15E-04 | 1.14E-04 | 5.404 | 3.27E-08 | 1.66E-03 |
| 3 | 85582231 | 87409731 | 3043 | 6.89E-04 | 1.17E-04 | 5.878 | 2.07E-09 | 1.86E-03 |
| 3 | 87409732 | 88298372 | 1607 | 4.08E-04 | 9.38E-05 | 4.349 | 6.84E-06 | 1.10E-03 |
| 3 | 113947863 | 115447079 | 1636 | 5.11E-04 | 1.09E-04 | 4.693 | 1.35E-06 | 1.38E-03 |
| 3 | 116800153 | 118530250 | 2960 | 7.85E-04 | 1.21E-04 | 6.466 | 5.03E-11 | 2.11E-03 |
| 3 | 126214943 | 128194860 | 3074 | 4.78E-04 | 1.07E-04 | 4.452 | 4.26E-06 | 1.29E-03 |
| 3 | 131836516 | 133252172 | 2767 | 4.96E-04 | 1.08E-04 | 4.582 | 2.30E-06 | 1.34E-03 |
| 3 | 185068255 | 186890343 | 3040 | 5.63E-04 | 1.11E-04 | 5.054 | 2.16E-07 | 1.52E-03 |
| 4 | 27965868 | 29762207 | 3857 | 4.24E-04 | 1.05E-04 | 4.051 | 2.55E-05 | 1.14E-03 |
| 4 | 43965045 | 45189156 | 2620 | 7.84E-04 | 1.21E-04 | 6.464 | 5.11E-11 | 2.11E-03 |
| 4 | 100678360 | 103221355 | 4515 | 4.77E-04 | 1.07E-04 | 4.444 | 4.41E-06 | 1.29E-03 |
| 4 | 103221356 | 105305293 | 2773 | 9.86E-04 | 1.30E-04 | 7.599 | 1.49E-14 | 2.66E-03 |
| 4 | 105305294 | 107501304 | 3048 | 4.24E-04 | 1.05E-04 | 4.048 | 2.58E-05 | 1.14E-03 |
| 5 | 41888710 | 43983498 | 2562 | 4.60E-04 | 1.06E-04 | 4.316 | 7.95E-06 | 1.24E-03 |
| 5 | 58524622 | 60935906 | 3047 | 4.87E-04 | 1.08E-04 | 4.515 | 3.16E-06 | 1.31E-03 |
| 5 | 75798866 | 77623331 | 3089 | 4.67E-04 | 1.07E-04 | 4.368 | 6.28E-06 | 1.26E-03 |
| 5 | 87389991 | 88891529 | 1261 | 5.90E-04 | 1.10E-04 | 5.351 | 4.39E-08 | 1.59E-03 |
| 5 | 93809984 | 95963518 | 2866 | 4.91E-04 | 1.08E-04 | 4.548 | 2.71E-06 | 1.32E-03 |
| 5 | 110821144 | 111981561 | 2081 | 7.76E-04 | 1.21E-04 | 6.412 | 7.20E-11 | 2.09E-03 |

| | | | | | | | | |
|---|---|---|---|---|---|---|---|---|
| 5 | 132139649 | 134777400 | 3641 | 7.61E-04 | 1.20E-04 | 6.324 | 1.27E-10 | 2.05E-03 |
| 5 | 136376050 | 139265071 | 2805 | 4.99E-04 | 1.08E-04 | 4.606 | 2.06E-06 | 1.35E-03 |
| 5 | 166847740 | 168525317 | 2749 | 5.74E-04 | 1.12E-04 | 5.125 | 1.49E-07 | 1.55E-03 |
| 5 | 180559933 | 180885155 | 259 | 3.08E-04 | 7.32E-05 | 4.205 | 1.31E-05 | 8.29E-04 |
| 6 | 28017819 | 28917607 | 1580 | 4.24E-04 | 8.81E-05 | 4.818 | 7.26E-07 | 1.14E-03 |
| 6 | 28917608 | 29737970 | 1999 | 4.53E-04 | 9.96E-05 | 4.553 | 2.64E-06 | 1.22E-03 |
| 6 | 30798168 | 31571217 | 5785 | 4.79E-04 | 1.07E-04 | 4.459 | 4.11E-06 | 1.29E-03 |
| 6 | 40345115 | 42038720 | 3015 | 5.54E-04 | 1.11E-04 | 4.992 | 2.98E-07 | 1.49E-03 |
| 6 | 50386145 | 52210475 | 3052 | 4.39E-04 | 1.05E-04 | 4.165 | 1.55E-05 | 1.18E-03 |
| 6 | 53279102 | 55468269 | 4522 | 5.77E-04 | 1.12E-04 | 5.146 | 1.33E-07 | 1.55E-03 |
| 6 | 56105313 | 57599464 | 1645 | 6.41E-04 | 1.15E-04 | 5.574 | 1.25E-08 | 1.73E-03 |
| 6 | 75462003 | 77414880 | 2539 | 5.24E-04 | 1.10E-04 | 4.784 | 8.59E-07 | 1.41E-03 |
| 6 | 77414881 | 78957727 | 3127 | 7.23E-04 | 1.19E-04 | 6.093 | 5.56E-10 | 1.95E-03 |
| 6 | 83127149 | 85209988 | 2529 | 4.29E-04 | 1.05E-04 | 4.083 | 2.22E-05 | 1.15E-03 |
| 6 | 97842284 | 100630145 | 4056 | 1.08E-03 | 1.34E-04 | 8.112 | 2.50E-16 | 2.92E-03 |
| 6 | 100630146 | 102636771 | 2725 | 1.10E-03 | 1.34E-04 | 8.220 | 1.02E-16 | 2.98E-03 |
| 6 | 103983395 | 106056732 | 3662 | 4.21E-03 | 2.24E-04 | 18.735 | 1.28E-78 | 1.13E-02 |
| 6 | 108464380 | 110304246 | 2480 | 4.39E-04 | 1.05E-04 | 4.161 | 1.58E-05 | 1.18E-03 |
| 6 | 125424383 | 127540460 | 2016 | 7.01E-04 | 1.18E-04 | 5.956 | 1.30E-09 | 1.89E-03 |
| 6 | 150253404 | 151912702 | 3899 | 4.54E-04 | 1.06E-04 | 4.272 | 9.71E-06 | 1.22E-03 |
| 7 | 1353067 | 2062397 | 1350 | 5.97E-04 | 1.13E-04 | 5.280 | 6.47E-08 | 1.61E-03 |
| 7 | 31137289 | 33555767 | 4922 | 4.44E-04 | 1.06E-04 | 4.197 | 1.35E-05 | 1.20E-03 |
| 7 | 39862670 | 42001810 | 2805 | 6.57E-04 | 1.16E-04 | 5.677 | 6.85E-09 | 1.77E-03 |
| 7 | 73334602 | 76458563 | 2899 | 6.35E-04 | 1.15E-04 | 5.533 | 1.58E-08 | 1.71E-03 |
| 7 | 121933630 | 124156804 | 3641 | 5.80E-04 | 1.12E-04 | 5.172 | 1.16E-07 | 1.56E-03 |
| 8 | 3392926 | 3783016 | 1951 | 5.58E-04 | 1.11E-04 | 5.021 | 2.57E-07 | 1.50E-03 |
| 8 | 4480476 | 5146926 | 2374 | 7.08E-04 | 1.18E-04 | 5.998 | 9.99E-10 | 1.91E-03 |
| 8 | 53302930 | 54418043 | 2183 | 8.89E-04 | 1.26E-04 | 7.069 | 7.82E-13 | 2.40E-03 |
| 8 | 76456542 | 79132860 | 4439 | 1.39E-03 | 1.45E-04 | 9.575 | 5.08E-22 | 3.74E-03 |
| 9 | 6557589 | 7154922 | 1628 | 5.76E-04 | 1.12E-04 | 5.144 | 1.34E-07 | 1.55E-03 |
| 9 | 9166403 | 10879252 | 4210 | 4.77E-04 | 1.07E-04 | 4.445 | 4.39E-06 | 1.29E-03 |
| 9 | 10879253 | 12276488 | 3463 | 4.86E-04 | 1.08E-04 | 4.509 | 3.25E-06 | 1.31E-03 |
| 9 | 12276489 | 14836362 | 5301 | 5.40E-04 | 1.10E-04 | 4.897 | 4.87E-07 | 1.46E-03 |
| 9 | 74253040 | 76203945 | 2929 | 4.47E-04 | 1.06E-04 | 4.220 | 1.22E-05 | 1.20E-03 |
| 9 | 76203946 | 76973080 | 1036 | 5.55E-04 | 9.93E-05 | 5.593 | 1.11E-08 | 1.50E-03 |
| 9 | 82590928 | 84211232 | 3771 | 4.31E-04 | 1.05E-04 | 4.106 | 2.02E-05 | 1.16E-03 |
| 9 | 85440801 | 86938195 | 2650 | 5.59E-04 | 1.11E-04 | 5.022 | 2.56E-07 | 1.51E-03 |
| 9 | 107581749 | 109298753 | 2824 | 2.20E-03 | 1.72E-04 | 12.796 | 8.68E-38 | 5.92E-03 |
| 9 | 126971887 | 129059664 | 2248 | 7.59E-04 | 1.20E-04 | 6.311 | 1.38E-10 | 2.05E-03 |
| 10 | 63341695 | 65794113 | 4108 | 5.73E-04 | 1.12E-04 | 5.123 | 1.50E-07 | 1.55E-03 |
| 10 | 96221243 | 97822356 | 3034 | 7.28E-04 | 1.19E-04 | 6.124 | 4.57E-10 | 1.96E-03 |
| 10 | 97822357 | 100241301 | 3639 | 8.59E-04 | 1.24E-04 | 6.897 | 2.65E-12 | 2.31E-03 |
| 10 | 116421406 | 119523933 | 4352 | 5.75E-04 | 1.12E-04 | 5.133 | 1.43E-07 | 1.55E-03 |
| 10 | 123231465 | 123900544 | 1414 | 4.23E-04 | 1.05E-04 | 4.044 | 2.63E-05 | 1.14E-03 |
| 10 | 125869346 | 128001097 | 4291 | 4.61E-04 | 1.07E-04 | 4.330 | 7.46E-06 | 1.24E-03 |

| | | | | | | | | |
|---|---|---|---|---|---|---|---|---|
| 11 | 8333274 | 9087316 | 1299 | 7.18E-04 | 1.18E-04 | 6.062 | 6.72E-10 | 1.94E-03 |
| 11 | 12564229 | 13373123 | 1662 | 8.55E-04 | 1.24E-04 | 6.876 | 3.08E-12 | 2.30E-03 |
| 11 | 15742552 | 17578401 | 2587 | 4.50E-04 | 1.06E-04 | 4.245 | 1.09E-05 | 1.21E-03 |
| 11 | 27020461 | 28481592 | 1698 | 7.67E-04 | 1.21E-04 | 6.360 | 1.01E-10 | 2.07E-03 |
| 11 | 28481593 | 30141356 | 2085 | 6.89E-04 | 1.17E-04 | 5.879 | 2.07E-09 | 1.86E-03 |
| 11 | 42310003 | 44693798 | 4484 | 4.31E-04 | 1.05E-04 | 4.100 | 2.07E-05 | 1.16E-03 |
| 11 | 44693799 | 47006136 | 2991 | 6.03E-04 | 1.13E-04 | 5.325 | 5.05E-08 | 1.63E-03 |
| 11 | 47006137 | 49866049 | 4586 | 6.97E-04 | 1.18E-04 | 5.932 | 1.50E-09 | 1.88E-03 |
| 11 | 55082657 | 58457494 | 7976 | 4.26E-04 | 1.05E-04 | 4.064 | 2.42E-05 | 1.15E-03 |
| 11 | 76797209 | 78355057 | 2236 | 4.88E-04 | 1.08E-04 | 4.523 | 3.04E-06 | 1.31E-03 |
| 11 | 101331121 | 103959635 | 5772 | 4.61E-04 | 1.07E-04 | 4.327 | 7.55E-06 | 1.24E-03 |
| 11 | 122591910 | 123500116 | 1750 | 1.18E-03 | 1.37E-04 | 8.604 | 3.86E-18 | 3.19E-03 |
| 12 | 23820634 | 25371082 | 2958 | 5.44E-04 | 1.11E-04 | 4.919 | 4.36E-07 | 1.46E-03 |
| 12 | 49001866 | 51776493 | 3252 | 7.10E-04 | 1.18E-04 | 6.014 | 9.04E-10 | 1.91E-03 |
| 12 | 97108839 | 99305986 | 3796 | 4.53E-04 | 1.06E-04 | 4.268 | 9.87E-06 | 1.22E-03 |
| 12 | 119754110 | 122007650 | 3707 | 4.54E-04 | 1.06E-04 | 4.276 | 9.53E-06 | 1.22E-03 |
| 13 | 27284362 | 29257550 | 4183 | 6.35E-04 | 1.15E-04 | 5.531 | 1.59E-08 | 1.71E-03 |
| 13 | 38878163 | 41069262 | 3744 | 5.85E-04 | 1.12E-04 | 5.199 | 1.00E-07 | 1.58E-03 |
| 13 | 61591949 | 63971558 | 4903 | 5.44E-04 | 1.11E-04 | 4.923 | 4.26E-07 | 1.47E-03 |
| 13 | 71526791 | 73934088 | 4070 | 4.67E-04 | 1.07E-04 | 4.372 | 6.15E-06 | 1.26E-03 |
| 13 | 73934089 | 75670142 | 3207 | 6.48E-04 | 1.15E-04 | 5.619 | 9.60E-09 | 1.75E-03 |
| 14 | 59448336 | 61680423 | 3132 | 6.21E-04 | 1.14E-04 | 5.444 | 2.61E-08 | 1.67E-03 |
| 14 | 99138532 | 101534306 | 3845 | 7.28E-04 | 1.19E-04 | 6.122 | 4.62E-10 | 1.96E-03 |
| 15 | 21131604 | 24195126 | 1976 | 7.61E-04 | 1.20E-04 | 6.325 | 1.26E-10 | 2.05E-03 |
| 15 | 24195127 | 25108589 | 2915 | 4.26E-04 | 1.05E-04 | 4.063 | 2.43E-05 | 1.15E-03 |
| 15 | 41177514 | 42776398 | 2313 | 5.69E-04 | 1.12E-04 | 5.096 | 1.74E-07 | 1.53E-03 |
| 15 | 59694116 | 61265835 | 2624 | 5.78E-04 | 1.12E-04 | 5.152 | 1.29E-07 | 1.56E-03 |
| 15 | 76398624 | 78516052 | 3060 | 4.20E-04 | 1.05E-04 | 4.022 | 2.89E-05 | 1.13E-03 |
| 15 | 80860978 | 84260467 | 3792 | 4.62E-04 | 1.07E-04 | 4.333 | 7.35E-06 | 1.24E-03 |
| 15 | 88369943 | 90475550 | 3283 | 6.29E-04 | 1.14E-04 | 5.492 | 1.99E-08 | 1.69E-03 |
| 16 | 2764829 | 4001195 | 1814 | 4.59E-04 | 1.06E-04 | 4.315 | 8.00E-06 | 1.24E-03 |
| 16 | 5891580 | 6893550 | 2949 | 4.38E-04 | 1.05E-04 | 4.158 | 1.60E-05 | 1.18E-03 |
| 16 | 13154437 | 14464001 | 2622 | 5.96E-04 | 1.13E-04 | 5.276 | 6.62E-08 | 1.61E-03 |
| 16 | 18643607 | 20150570 | 2314 | 4.94E-04 | 1.08E-04 | 4.571 | 2.43E-06 | 1.33E-03 |
| 16 | 29036613 | 31382942 | 1785 | 5.01E-04 | 1.08E-04 | 4.615 | 1.96E-06 | 1.35E-03 |
| 16 | 53382572 | 55903773 | 4860 | 1.03E-03 | 1.31E-04 | 7.825 | 2.55E-15 | 2.77E-03 |
| 16 | 68841363 | 71054027 | 2201 | 9.84E-04 | 1.30E-04 | 7.593 | 1.57E-14 | 2.65E-03 |
| 17 | 43056905 | 45876021 | 4503 | 6.60E-04 | 1.16E-04 | 5.693 | 6.24E-09 | 1.78E-03 |
| 17 | 45876022 | 47517399 | 2620 | 4.21E-04 | 1.05E-04 | 4.024 | 2.86E-05 | 1.13E-03 |
| 17 | 51826118 | 53599431 | 2958 | 5.78E-04 | 1.12E-04 | 5.152 | 1.29E-07 | 1.56E-03 |
| 17 | 55357541 | 57487511 | 2720 | 5.14E-04 | 1.09E-04 | 4.709 | 1.25E-06 | 1.38E-03 |
| 17 | 64800430 | 67858769 | 4759 | 4.74E-04 | 1.07E-04 | 4.423 | 4.86E-06 | 1.28E-03 |
| 17 | 78837588 | 80034407 | 2477 | 4.43E-04 | 1.06E-04 | 4.192 | 1.38E-05 | 1.19E-03 |
| 18 | 31780067 | 33861963 | 2881 | 4.26E-04 | 1.05E-04 | 4.063 | 2.43E-05 | 1.15E-03 |
| 18 | 44299246 | 45939731 | 2694 | 9.59E-04 | 1.29E-04 | 7.457 | 4.44E-14 | 2.59E-03 |

| | | | | | | | | |
|---|---|---|---|---|---|---|---|---|
| 18 | 51554175 | 55213837 | 4947 | 4.31E-04 | 1.05E-04 | 4.098 | 2.08E-05 | 1.16E-03 |
| 18 | 57630483 | 59020750 | 2079 | 4.36E-04 | 1.05E-04 | 4.137 | 1.76E-05 | 1.17E-03 |
| 19 | 610729 | 2098395 | 2998 | 7.12E-04 | 1.18E-04 | 6.022 | 8.61E-10 | 1.92E-03 |
| 19 | 9238393 | 11284027 | 3512 | 7.87E-04 | 1.21E-04 | 6.478 | 4.64E-11 | 2.12E-03 |
| 19 | 18409862 | 19877470 | 1993 | 5.12E-04 | 1.09E-04 | 4.699 | 1.31E-06 | 1.38E-03 |
| 19 | 34262952 | 36469294 | 3728 | 4.25E-04 | 1.05E-04 | 4.058 | 2.48E-05 | 1.15E-03 |
| 19 | 47150082 | 49282226 | 3562 | 6.49E-04 | 1.15E-04 | 5.622 | 9.44E-09 | 1.75E-03 |
| 20 | 8117011 | 9730920 | 2887 | 4.25E-04 | 1.05E-04 | 4.060 | 2.45E-05 | 1.15E-03 |
| 20 | 15958359 | 17487231 | 3557 | 5.74E-04 | 1.12E-04 | 5.125 | 1.49E-07 | 1.55E-03 |
| 20 | 18972203 | 20959709 | 3062 | 4.23E-04 | 1.05E-04 | 4.043 | 2.64E-05 | 1.14E-03 |
| 20 | 32813441 | 34960445 | 2123 | 5.21E-04 | 1.09E-04 | 4.759 | 9.71E-07 | 1.40E-03 |
| 21 | 36524373 | 37870778 | 2570 | 4.42E-04 | 1.06E-04 | 4.186 | 1.42E-05 | 1.19E-03 |
| 21 | 40482902 | 41389526 | 2361 | 5.19E-04 | 1.09E-04 | 4.744 | 1.05E-06 | 1.40E-03 |
| 22 | 29651799 | 31439917 | 2535 | 4.32E-04 | 1.05E-04 | 4.106 | 2.01E-05 | 1.16E-03 |
| **c) Age at first birth** | | | | | | | | |
| 3 | 47727212 | 49316971 | 1015 | 4.46E-04 | 7.06E-05 | 6.315 | 1.35E-10 | 7.37E-03 |
| 3 | 49316972 | 51832014 | 2127 | 4.68E-04 | 7.85E-05 | 5.959 | 1.27E-09 | 7.74E-03 |

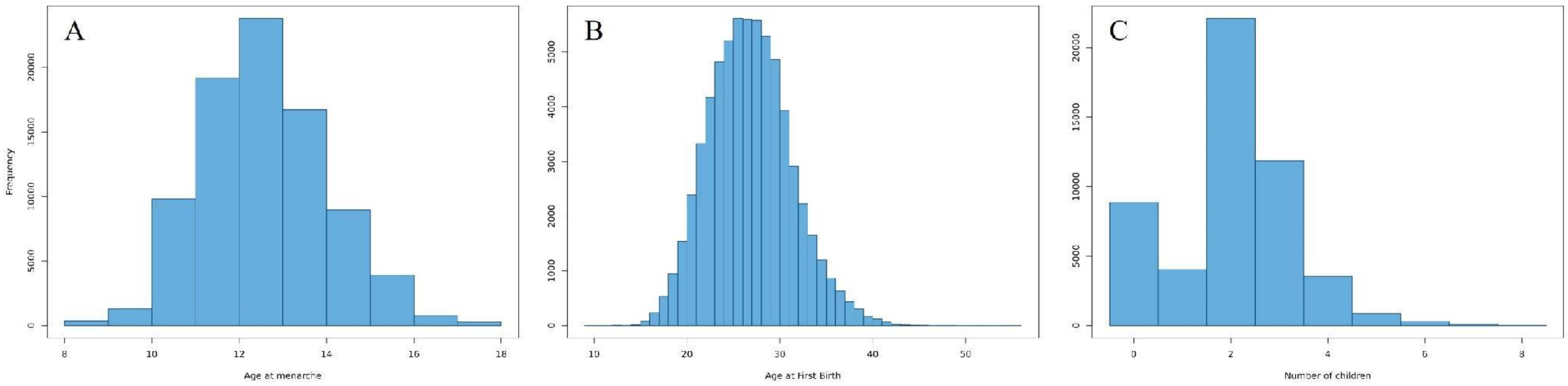

Figure S4: Histograms of life-history traits: (**A**) age at menarche, (**B**) age at first birth, and (**C**) number of children.

### SUPPLEMENTARY NOTE 1

We subtyped these records according to oestrogen receptor status (ER-status) using the report conclusions written by pathologists contained in the PALGA records. According to national Dutch guidelines, ER-status is considered positive when >= 10% of tumour cells show ER-specific staining, and otherwise ER-negative, when using immunohistochemistry analysis of tumour tissue of formalin-fixed and paraffin-embedded tumour material (IKNL, 2012). Subtyping rates across Dutch laboratories show limited variation suggesting strong standardisation (Van Dooijeweert et al., 2019).

If there were multiple contradictory ER-statuses within the same report conclusions (n = 20), the following rules applied: (1) if statuses differed between a biopsy and resection from the same tumour, results from the more reliable resections were used (n = 9); (2) if the conclusion contained a follow-up note correcting the ER-status, the correction was followed (n = 2); and (3) if there were differing statuses across multiple tumours, the status from the larger or original tumour was used (n = 1). In cases of multiple tumours with different ER-statuses and no clear origin, the ER-status of the tumour at that specific moment in time was set to unknown (n = 8). 4,769 individual reports (47.2%) did not include an ER-subtype assessment but mostly this was repeated reports on the same tumor overtime where the ER-status had already been assessed and was not needed to be reassessed.

We focused on women whose first cancer diagnosis was breast cancer, excluding 628 diagnoses to remove possible metastatic tumors of different origin. We also excluded 15 male diagnoses. This left 3,667 Lifelines participants with their first-cancer diagnosis being breast cancer, diagnosed 1981-2023, with a median age of diagnosis of 51 (range = 13-94). If the ER-status was not included in the first tumor report (n = 2,127), we classified ER status using reports within 3 months of the original diagnosis (ER+ = 849, ER- = 189), as ER-status is often accessed by pathologists in follow-up analyses after initial diagnoses are made. Extrapolating records beyond three months risked incorrectly subtyping the tumor because ER-status of tumors can change overtime in response to treatment. We also checked for inconsistencies between the ER-statuses of individual tumors over time. As before, we used the ER-statuses from resections over biopsies when inconsistent (n = 6) and changed the ER-status of 1 tumor to positive after a follow-up analysis was used to confirm an inconclusive result. However, we excluded bilateral tumors where ER-status differed (n = 9). In total, this resulted in the assignment of 2,177 ER+ first-cancer diagnoses (Figure S5) and 415 ER- diagnoses, with 1,090 being unable to be assigned an ER-status. In line with previous literature, ER+ cases showed a later age of onset than ER- cases (median 53 vs 49) (Yasui & Potter, 1999).

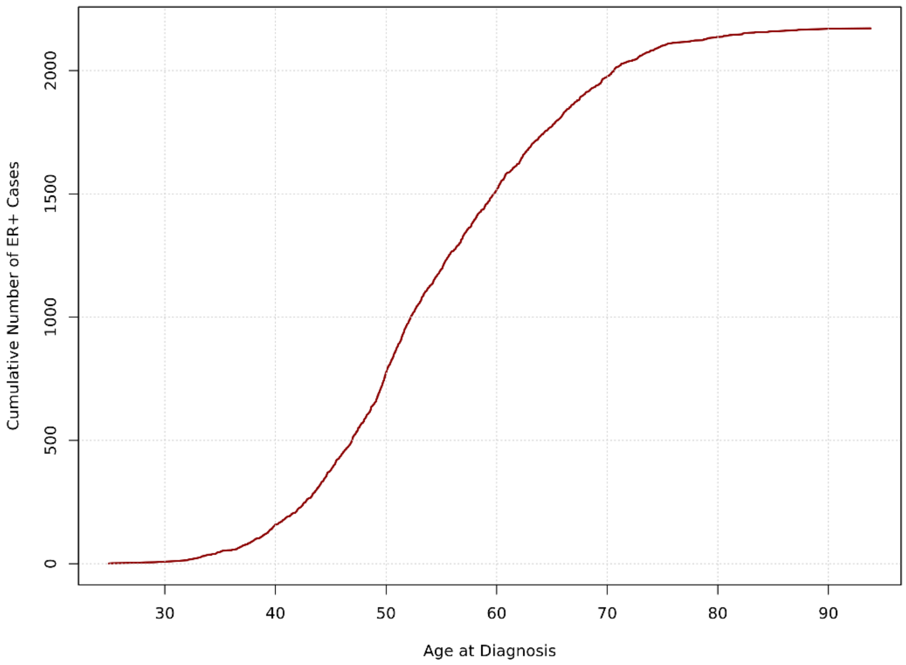

Figure S5: Density plot showing the cumulative number of ER+ breast cancer cases by age at diagnosis across Lifelines individuals in our data.
